# The gp120 Envelope Glycoprotein of HIV-1 Triggers Macropinocytosis in Primary CD4+ T Cells to Promote HIV-1 Infection

**DOI:** 10.64898/2026.04.27.721102

**Authors:** Praveen Manivannan, Cristina Santos da Costa, Yipei Tang, Xinghao Wang, Joel Swanson, Tomoyuki Murakami, Akira Ono, Philip D. King

## Abstract

Macropinocytosis is a large-scale, actin-driven, fluid-phase form of endocytosis that can be exploited by HIV-1 for entry into primary CD4+ T cells. Herein, we show that HIV-1 actively induces macropinocytosis in resting and activated primary CD4+ T cells. HIV-1-induced T cell macropinocytosis is triggered by the interaction of soluble gp120 Env with cell surface CD4, is clathrin-independent and actin-dependent, and does not require CXCR4 or any other known HIV-1 coreceptors. Notably, Env-induced macropinocytosis requires calcium-independent phospholipase A2 (iPLA2) activity, identifying a role for a lipid-signaling axis downstream of Env–CD4 engagement. Across a gp120 panel, multiple HIV-1 clade B gp120 Envs can induce CD4+ T cell macropinocytosis. However, the magnitude of macropinocytosis does not correlate with CD4-binding strength alone and instead aligns with Env-driven changes in CD4 epitope exposure in CD4 domains distal to the Env-binding domain. Importantly, inhibitors of Env-induced macropinocytosis block HIV-1 infection of resting CD4+ T cells. Therefore, Env-induced macropinocytosis is a functionally important mechanism that promotes HIV-1 infection of this cell type.

## Introduction

Human immunodeficiency virus-1 (HIV-1) primarily infects peripheral CD4+ T cells and myeloid lineage cells of the human immune system, thereby causing acquired immunodeficiency syndrome (AIDS)^1,2^. HIV-1 infection of these cell types is initiated by the interaction of the viral surface trimeric envelope glycoprotein (Env) with the most membrane-distal immunoglobulin-like extracellular domain (D1) of the transmembrane CD4 protein^3,4^. This interaction triggers conformational changes in the Env trimer that expose the Env variable region loop 3 (V3 loop) and lead to the formation of the four-stranded Env bridging sheet^5–8^. Subsequently, the V3 loop and bridging sheet bind to the extracellular domains of the chemokine receptors CXCR4 or CCR5, thereby promoting the interaction of the Env fusion domain in the gp41 subunit of the trimer with the host cell membrane, leading to viral and host membrane fusion^5–8^.

While many details of the HIV-1 fusion reaction have been determined, the exact cellular location of viral fusion in primary cell targets has been slower to resolve. It is recognized that intact HIV-1 virions can be internalized into primary human T cells, monocytes and macrophages, and other cell types via endocytosis, raising the possibility that fusion could occur within endosomes as well as at the plasma membrane^9–14^. Certainly, detailed imaging analyses of primary CD4+ T cells and myeloid cells have supported the view that endosomes are a significant, if not the most significant, site of HIV-1 viral fusion^10–12^. Conversely, a study that examined the ability of the membrane-impermeable T-20 fusion inhibitor to block productive HIV-1 infection of primary CD4+ T cells, following internalization of virions into endosomes at 22°C, concluded that any HIV-1 fusion within endosomes did not significantly contribute to infection^15^. However, whereas clathrin-mediated endocytosis (CME) was not affected at 22°C in primary CD4+ T cells in these experiments, the effect of temperature on other forms of endocytosis was not examined.

Macropinocytosis is a large-scale fluid-phase form of endocytosis that is driven by actin cytoskeletal rearrangements, leading to the formation of membrane ruffles, macropinocytic cups, and eventually macropinosomes that can range from 0.1 to 1.0 µm in diameter^16–18^. Monocytes, macrophages, and dendritic cells actively engage in macropinocytosis, which they use to acquire extracellular antigens for processing and presentation to T cells^19,20^. In addition, Ras-transformed tumor cells employ macropinocytosis to acquire extracellular proteins to fuel cell growth and proliferation under amino acid-depleted conditions^21,22^. Significantly, HIV-1 is internalized into monocytes and macrophages and other cell types via macropinocytosis^9,11,13,14^. Moreover, macropinocytosis has been shown to mediate the uptake of a variety of other viruses into target cells, including influenza A, Ebola, vaccinia, and SARS-CoV-2^23–27^.

As relatively small cells with a paucity of cytoplasm, it was considered that primary T cells would not be able to perform macropinocytosis. However, we recently demonstrated that murine and human T cells do, in fact, engage in macropinocytosis that increases in magnitude upon T cell activation^28,29^. T cells use macropinocytosis to acquire extracellular amino acids, such as leucine and glutamine, which are delivered to lysosomes, where they activate the mechanistic target of rapamycin complex 1 (mTORC), which drives cell cycle progression^28,29^. In light of this discovery, we asked if HIV-1 might exploit macropinocytosis to infect primary human T cells. We found that HIV-1 was taken up via macropinocytosis in activated human CD4+ T cells and that HIV-1 fusion with the host cell membrane occurred at this site^30^. Furthermore, 5-(N-Ethyl-N-isopropyl) amiloride (EIPA), a highly specific inhibitor of macropinocytosis^31^, blocked HIV-1 entry into the cytoplasm of activated CD4+ T cells and inhibited productive viral infection^30^.

Given the important role of macropinocytosis in HIV-1 infection of primary CD4+ T cells, it is of interest to determine whether HIV-1 manipulates macropinocytosis to facilitate its infection of this cell type. In this regard, we observed that HIV-1 itself triggers macropinocytosis in both resting and activated primary human CD4+ T cells through a mechanism that involves soluble Env interaction with CD4 but is independent of chemokine co-receptors. Importantly, we show that HIV-1 Env-induced macropinocytosis contributes to HIV-1 infection of resting CD4+ T cells.

## Results

### HIV-1 triggers CD4+ T cell macropinocytosis

To investigate if HIV-1 could induce T cell macropinocytosis, peripheral blood mononuclear cells (PBMC) from healthy human donors were cultured with fluorescently labeled bovine serum albumin (BSA) in the presence of NL4-3 strain virus (clade B, CXCR4 tropic) produced by transient transfection of 293T cells. BSA is internalized into mammalian cells, including activated mouse and human T cells, predominantly via macropinocytosis and thus serves as a useful macropinocytosis probe^21,28,30,32^. As shown by flow cytometry after 2 h of culture, NL4-3 induced uptake of BSA into resting CD4+ T cells but not resting CD8+ T cells (Fig. 1a). Microscopic analyses of purified resting PB T cells cultured with NL4-3 in the presence of fluorescent BSA, confirmed that the probe was taken up into large macropinosome-like vesicles into CD4+ T cells but not CD8+ T cells (Fig. 1b). Furthermore, as shown by immunostaining for the HIV-1 p24 capsid protein, NL4-3 virions were frequently detected in the same vesicles as the BSA probe, consistent with the notion that HIV-1 can trigger its own internalization into resting CD4+ T cells via macropinocytosis (Fig. 1c).

**Figure 1.**
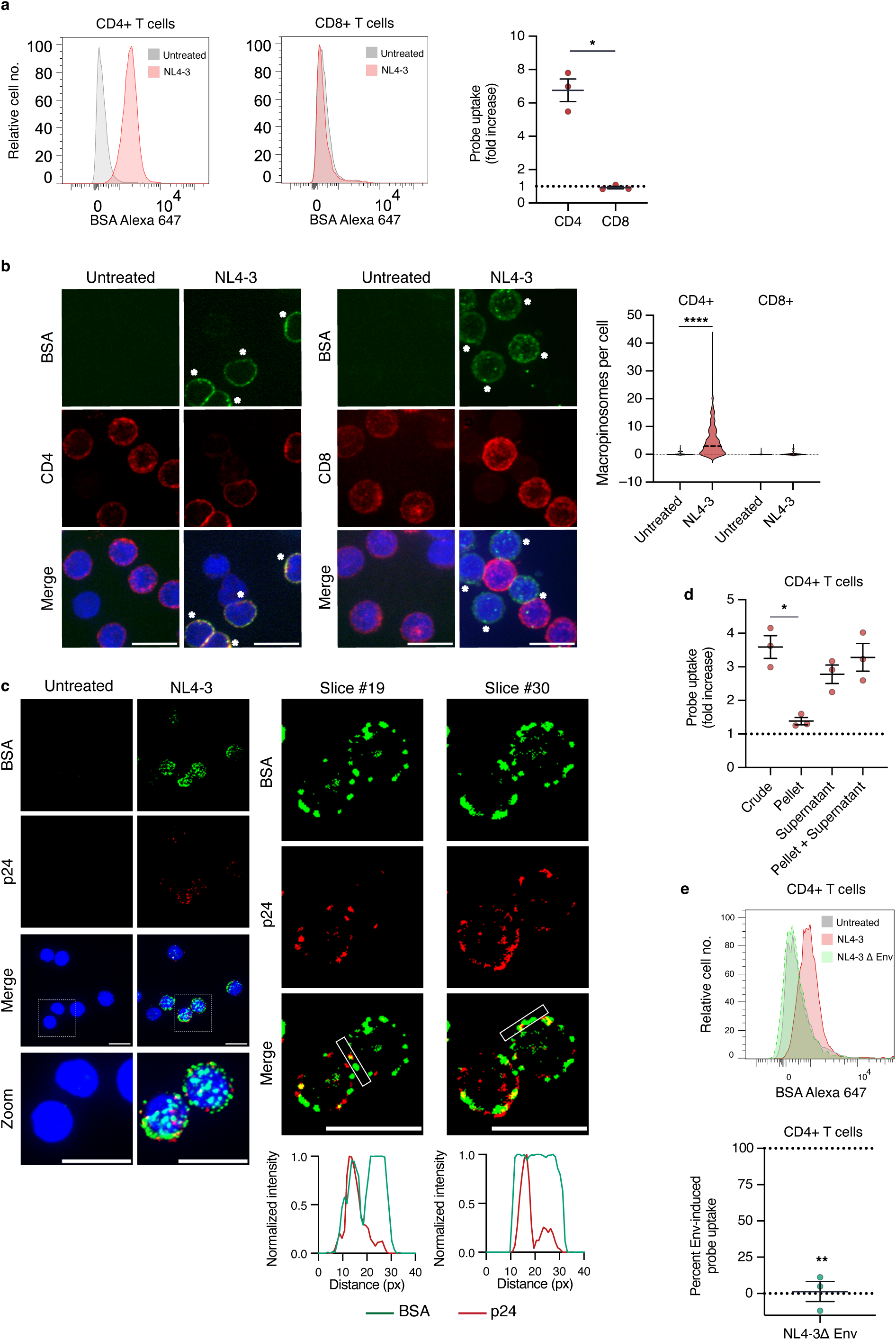
HIV-1 triggers macropinocytosis in resting CD4+ T cells. **(a)** PBMC from healthy donors were cultured with BSA-Alexa 647 in the presence of supernatants from 293T cells transiently transfected with HIV-1 molecular clone NL4-3 (clade B, CXCR4 tropic) for 2 h at 37°C. BSA uptake into gated CD4+ and CD8+ T cell populations was determined by flow cytometry. Representative flow cytometry histograms are shown at left. The graph at right shows the fold increase in probe uptake in HIV-1-treated versus untreated T cells, based on median fluorescence intensity (MFI), from independent donors. Mean ± 1 SEM fold increases in probe uptake are depicted. Statistical significance was determined using a Mann-Whitney *U* test. **(b)** Microscopic analysis of resting PB T cells cultured or not with NL4-3 virus-containing supernatants in the presence of BSA-Alexa 647 for 2 h. Cells were counterstained with anti-CD4 or anti-CD8 antibodies to distinguish CD4+ and CD8+ T cells. Left, Maximal Intensity Projection (MIP) images are shown. Asterisks indicate CD4+ T cells. Scale bar = 10 µm. Right, violin plot showing the number of macropinosomes per cell (n=115, 368, 81, and 50 for CD4+ untreated and NL4-3 treated, and CD8+ untreated and NL4-3 treated, respectively). Statistical significance was determined using a Kruskal-Wallis test with Dunn’s correction. **(c)** Purified resting CD4+ T cells cultured or not with NL4-3 virus-containing supernatants in the presence of BSA-Alexa 647 for 2h were counterstained with an antibody against HIV-1 p24 capsid protein. MIP images are shown at left. Images on the right show select single slices of the NL4-3-treated samples to identify points of colocalization of BSA and p24. Graphs below show the intensity of green BSA and red p24 signals within the indicated windows in respective slices. Scale bar = 10 µm. **(d)** Fold induction of BSA uptake in resting CD4+ T cells within PBMC incubated with NL4-3 virus-containing 293T supernatants (crude), NL4-3 virus purified by ultracentrifugation through sucrose (pellet), virus-depleted supernatants (supernatant), or pellet plus supernatant. Probe uptake was determined by flow cytometry. Results from independent donors are depicted. Statistical significance was determined using a Kruskal-Wallis test with Dunn’s correction, comparing each sample type to the crude virus as a reference. **(e)** The flow cytometry histogram shows BSA uptake by resting CD4+ T cells within PBMC in the presence of supernatants from 293T cells transfected with wild-type NL4-3 or NL4-3 lacking the Env glycoprotein. The graph shows the percent of probe uptake by NL4-3 lacking Env compared to wild-type NL4-3 from independent donors. Statistical significance was determined using a Student’s 1-sample t-test.

To determine if HIV-1 virions themselves induce CD4+ T cell BSA probe uptake, intact virus was purified from supernatants of transfected 293T cells by ultracentrifugation through sucrose. Surprisingly, although NL4-3 virus-containing 293T supernatants induced CD4+ T cell probe uptake, NL4-3 virus purified from these supernatants was much less active (Fig. 1d). In contrast, the same supernatants depleted of virus by ultracentrifugation retained activity, albeit reduced compared to non-spun 293T supernatants (Fig. 1d). Remixed pelleted virus and post-spin supernatant showed similar activity to that of non-spun 293T supernatants (Fig. 1d). These findings indicated that a soluble factor derived from HIV-1 and not intact HIV-1 virions themselves were mostly responsible for CD4+ T cell macropinocytosis. A strong candidate is the gp120 subunit of the gp160 HIV-1 envelope glycoprotein (Env), which is known to be shed from HIV-1 virions during virus production in vitro and during HIV-1 infection in vivo^33–37^. In agreement with this possibility, supernatants containing NL4-3 virus that lacked the Env glycoprotein were unable to induce BSA probe uptake in resting CD4+ T cells (Fig. 1e).

### Soluble HIV-1 Env induces CD4+ T cell macropinocytosis

To directly address whether shed Env was responsible for the induction of macropinocytosis by HIV-1-containing viral supernatants, we examined whether soluble recombinant gp120 HIV-1 Env could induce macropinocytosis in CD4+ T cells. As shown by flow cytometry, recombinant soluble gp120 from HIV-1 strains IIIB (clade B, CXCR4 tropic) and BAL (clade B, CXCR5 tropic) induced uptake of BSA probe into resting CD4+ T cells but not resting CD8+ T cells in mixed PBMC populations and purified resting CD4+ T cells (Fig. 2a, b). The extent of probe uptake depended on Env concentration and exposure duration, with CD4+ T cells reaching a plateau in fluorescence after 4 h of Env stimulation (Fig. 2c, d). Importantly, Env-induced probe uptake was observed at 37°C, but not 4°C, consistent with an active energy-dependent process (Fig. 2e). Furthermore, probe uptake was much reduced at 22°C compared to 37°C, consistent with the notion that T cell macropinocytosis is restricted at the former temperature, which, in turn, provides an explanation for earlier conflicting findings regarding the role of endocytosis in HIV-1 infection of primary CD4+ T cells^10,12,15,30^ (Fig. 2e). Microscopic analyses confirmed that soluble gp120 HIV-1 Env induced the uptake of probe into macropinosome-like structures in resting CD4+ T cells (Fig. 2f).

**Figure 2.**
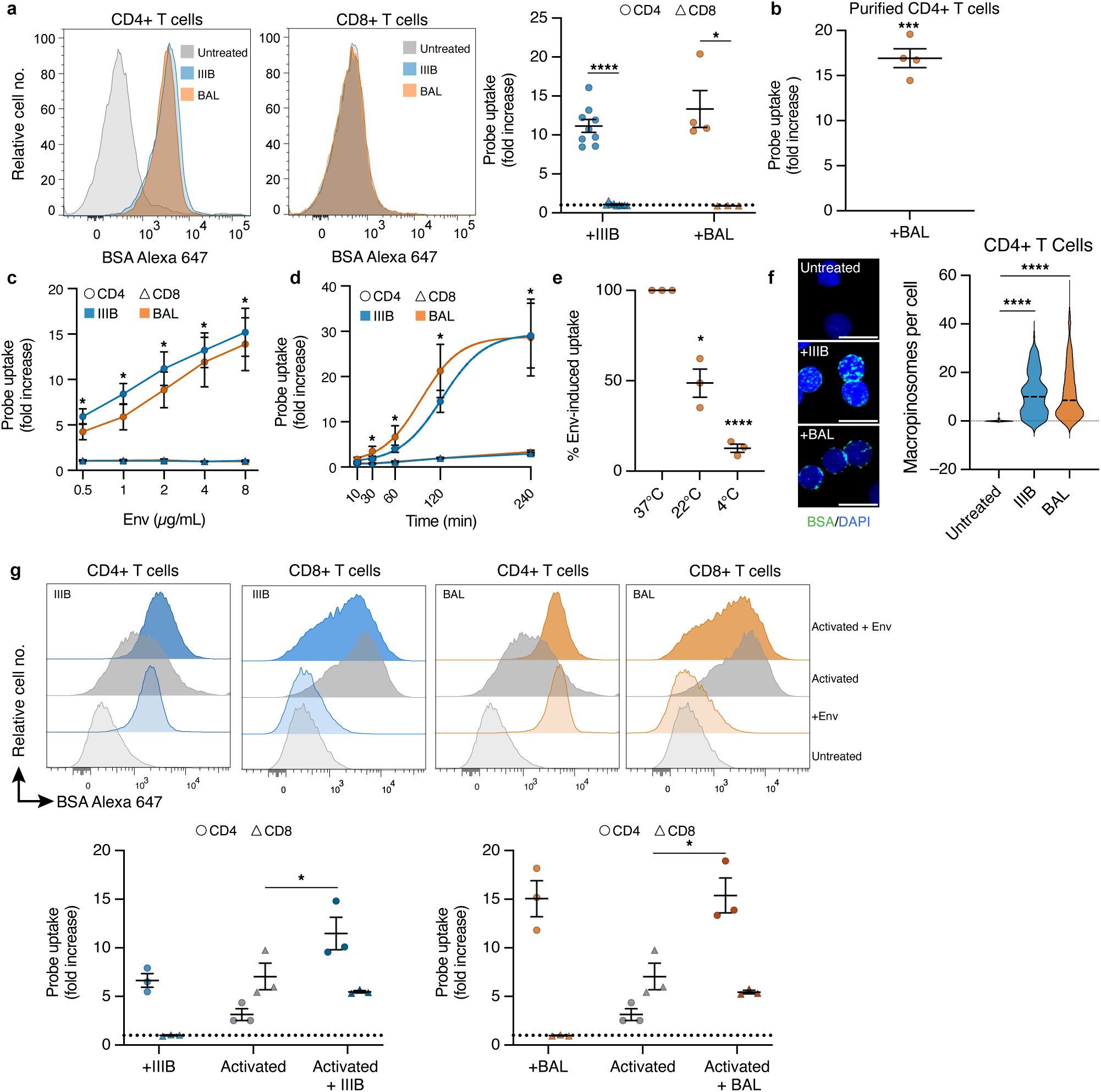
Soluble gp120 HIV-1 Env induces CD4+ T cell macropinocytosis. (a-e) PBMC (a, c-e) or purified CD4+ T cells (b) were incubated with BSA-Alexa 647 and soluble recombinant gp120 from HIV-1 strains IIIB (clade B, CXCR4 tropic) or BAL (clade B, CCR5 tropic) at 2 µg/ml (a,b,d,e) or at the concentrations indicated (c) for 2 h at 37°C (a-c), or for the time indicated at 37°C (d), or for 2 h at the temperatures indicated (e). Flow cytometry histograms show representative probe uptake in CD4+ and CD8+ T cells (a). Graphs show fold increases in BSA uptake in CD4+ and CD8+ T cells from independent donors. Statistical significance was determined using a Mann-Whitney *U* test (a,c,d) or Student’s 1-sample t-test (b,e). **(f)** Microscopic analysis of resting CD4+ T cells following treatment with soluble HIV-1 Env in the presence of BSA-Alexa 647. Representative images are shown on the left; quantitation is shown on the right (n= 27, 129, 212 for untreated, IIIB, and BAL, respectively). Statistical significance was determined using a Kruskal-Wallis test with Dunn’s correction. **(g)** Resting and PHA-activated PBMC were incubated with gp120 IIIB or BAL (2 µg/ml) in the presence of BSA-Alexa 647 for 2 h at 37°C. Representative histograms showing probe uptake in CD4+ and CD8+ T cells are shown at top. Graphs depicting fold increase in BSA uptake relative to unactivated CD4+ and CD8+ T cells from independent donors are shown at the bottom. Statistical significance was determined using a Mann-Whitney *U* test.

HIV-1 virion-associated Env exists as a trimer of three Env protomers, with each protomer comprising a gp120 subunit non-covalently linked to a gp41 subunit^38^. We considered the possibility that trimeric Env might be much less able to induce macropinocytosis in CD4+ T cells, thereby explaining the relative inability of intact HIV-1 virions to trigger this event. To test this possibility, we examined the ability of soluble gp140 recombinant trimeric Env from HIV-1 strain SF162 (clade B, CCR5 tropic) to induce CD4+ T cell macropinocytosis. The gp140 subunits in the SF162 trimer comprise gp120 covalently linked to a truncated gp41 to permit Env secretion. Trimeric gp140 SF162 Env was able to induce CD4+ T cell macropinocytosis (Supplementary Figure 1). Therefore, the relative inability of intact HIV-1 virions to induce T cell macropinocytosis in our experiments is unlikely to be explained by the trimeric nature of virion-associated Env.

Resting human CD4+ and CD8+ T cells endocytose minimal amounts of BSA probe^28^. However, as referenced above, activation of T cells, using TCR/CD28 agonists or phytohemagglutinin (PHA) for 24 h stimulates probe uptake by several fold^28^. Therefore, we asked if soluble HIV-1 Env could augment probe uptake of pre-activated T cells that would be potentially relevant to HIV-1 infection of activated CD4+ T cells in vivo. Indeed, soluble HIV-1 Env stimulated activated CD4+ T cells to endocytose increased amounts of probe (Fig. 2g). In contrast, soluble HIV-1 Env did not augment probe uptake by activated CD8+ T cells. Therefore, the ability of HIV-1 Env to drive T cell macropinocytosis is not limited to quiescent CD4+ T cells.

To confirm that soluble HIV-1 Env induced the internalization of BSA probe via macropinocytosis, we examined the effect of established macropinocytosis inhibitors on probe uptake. Macropinocytosis is driven by Rho-family GTPase-mediated actin polymerization, which leads to membrane ruffling and the formation of macropinocytic cups^16–18^. EIPA is a highly specific inhibitor of macropinocytosis that acts by blocking a plasma membrane Na^+^/H^+^ exchanger known as Nhe1^31^. Nhe1 functions in macropinocytosis by mitigating a submembrane drop in pH that would otherwise inhibit the activation of Rho GTPases. EIPA partially inhibited HIV-1 Env-induced probe uptake in resting CD4+ T cells (Fig. 3a). Similarly, the combination of the drugs Jasplakinolide and Blebbistatin (J/B), which impair actin cytoskeletal dynamics^39^, and the drugs EHT1864^40^ and ML141^41^ (E/M), which inhibit the Rac and Cdc42 Rho proteins, respectively, inhibited Env-induced probe uptake (Fig. 3a). In contrast, PitStop2, an inhibitor of clathrin-mediated endocytosis^42^, did not affect Env-induced probe uptake by CD4+ T cells (Fig. 3a).

**Figure 3.**
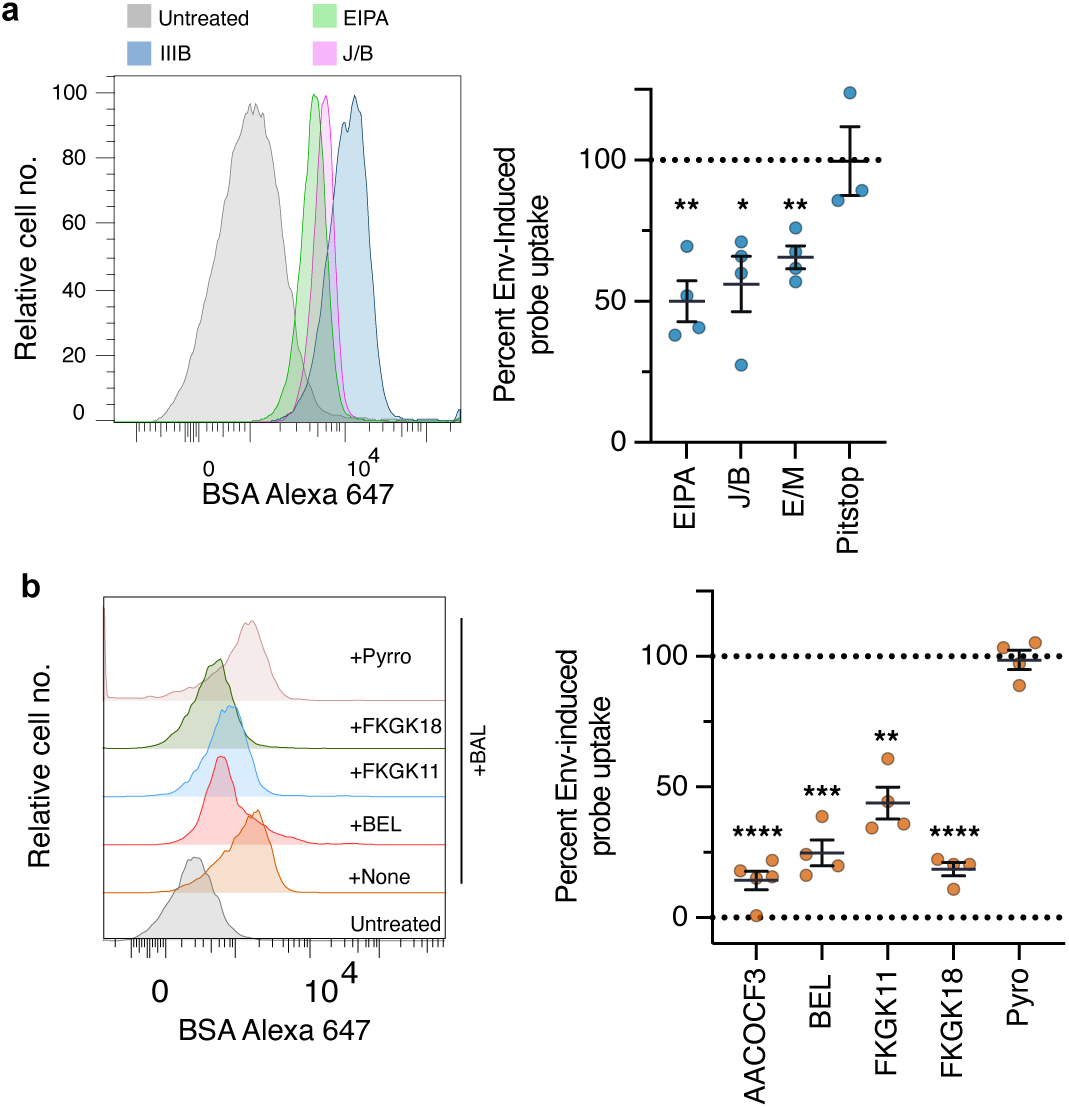
Env-induced probe uptake in CD4+ T cells is mediated predominantly by macropinocytosis. (a,b) PBMC were incubated with 2 µg/ml gp120 IIIB (a) or BAL (b) and BSA-Alexa 647 for 2 h at 37°C in the presence of the indicated endocytosis inhibitors (a) or PLA-2 inhibitors (b). Representative flow cytometry histograms showing probe uptake by CD4+ T cells are shown at left. Graphs on the right show the percent probe uptake by CD4+ T cells in the presence of inhibitors compared to the no-inhibitor control from independent donors. Statistical significance was determined using a Student’s 1-sample t-test.

Since EIPA, J/B, and E/M did not completely inhibit probe uptake, we examined other potential macropinocytosis inhibitors for their effects on Env-induced macropinocytosis. We established that arachidonyl trifluoromethyl ketone (AACOCF3)^43^, which targets cytosolic phospholipase A2 (cPLA-2) and calcium-independent PLA-2 (iPLA-2), potently inhibited PB monocyte macropinocytosis of BSA (Supplementary Fig. 2a). Further experiments revealed that more specific inhibitors of iPLA-2, including bromoenol lactone (BEL)^43^, and the fluoroketone-based compounds FKGK11^44^ and FKGK18^44,45^ were also highly active inhibitors of monocyte macropinocytosis (Supplementary Fig. 2a). In contrast, pyrrophenone (Pyro)^46^, a specific inhibitor of cPLA-2 had a minor effect (Supplementary Fig. 2a), thus pointing to an essential function iPLA-2 rather than cPLA-2 in monocyte macropinocytosis. Notably, AACOCF3 and FKGK18 also potently inhibited BSA probe uptake by PHA-activated human CD4+ and CD8+ T cells, consistent with an essential function for iPLA-2 in T cell macropinocytosis (Supplementary Fig. 2b). When tested for their effects on Env-induced probe uptake by resting CD4+ T cells, AACOCF3, BEL, and FKGK18 were also potent inhibitors. FKGK11 was also inhibitory, whereas Pyro was without effect (Fig. 3 b). These findings confirm that Env-induced probe uptake by CD4+ T cells is mediated predominantly by macropinocytosis.

### Env-induced T cell macropinocytosis requires Env interaction with CD4

The finding that HIV-1 Env induces macropinocytosis in CD4+ T cells but not CD8+ T cells suggests that Env binding to CD4 is required for this event. To confirm the role of Env-CD4 binding, we examined the effects of inhibitors of this interaction. A soluble fusion protein comprising the most N-terminal immunoglobulin (Ig)-like domain of CD4 (domain 1, D1) that contains the Env interaction site, fused to the constant region of human IgG1 (CD4Ig), completely blocked soluble gp120 Env-induced CD4+ T cell macropinocytosis (Fig. 4a). Likewise, a soluble protein consisting of all four extracellular Ig-like domains of CD4 (sCD4) completely blocked Env-induced macropinocytosis (Fig. 4a). Furthermore, all of three different examined monoclonal antibodies that bind epitopes on Env that block interaction with CD4 (N6, G54W, 65430D) completely blocked Env-induced macropinocytosis (Fig. 4b). CD4Ig, sCD4, and anti-Env antibodies that block Env interaction with CD4 also completely blocked NL4-3 viral supernatant-induced CD4+ T cell macropinocytosis (Fig. 4a, b). Thus, Env-induced macropinocytosis in CD4+ T cells is dependent on Env physical interaction with CD4.

**Figure 4.**
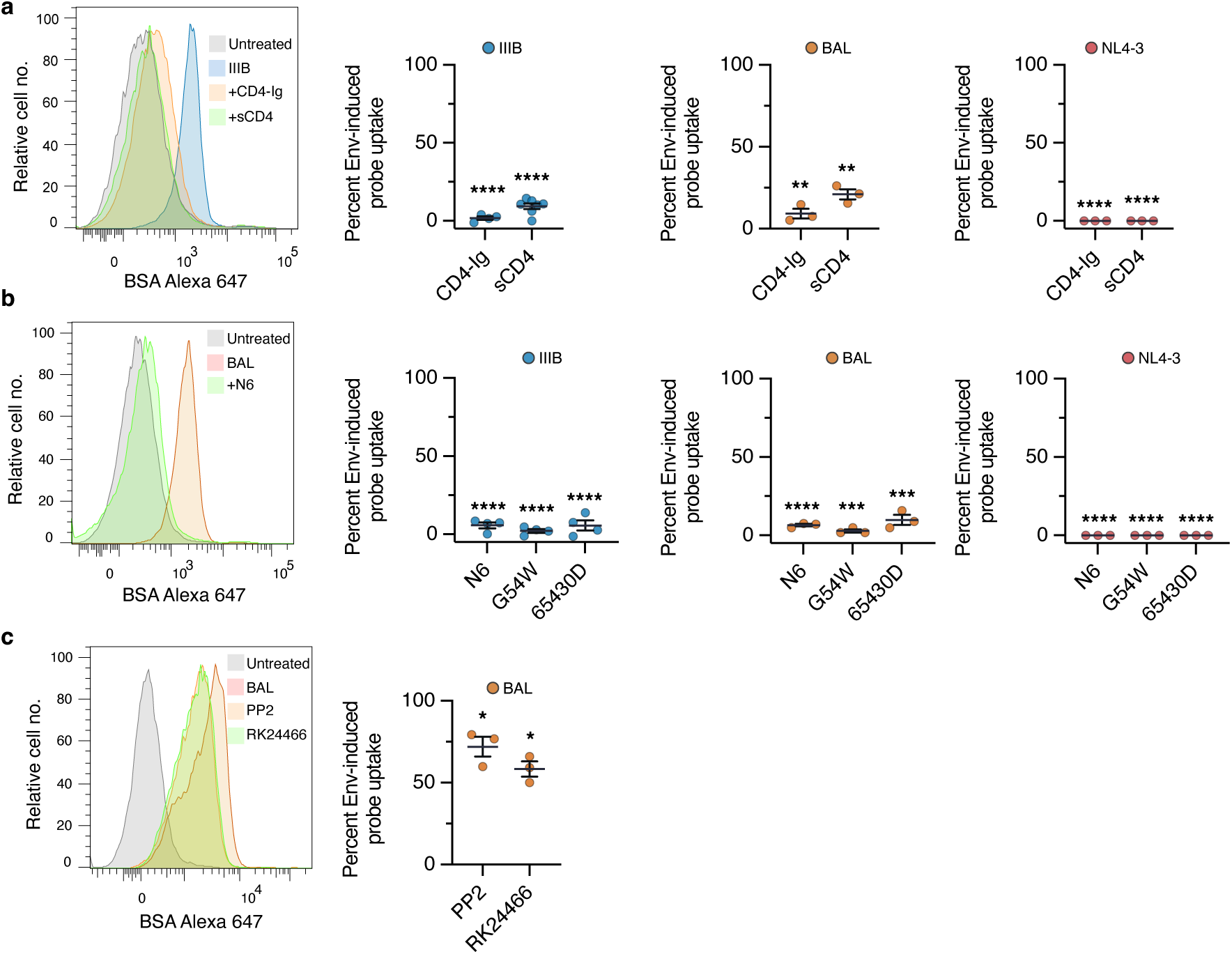
Env interaction with CD4 is required for Env-induced macropinocytosis. (a-c) PBMC were incubated with 2 µg/ml gp120 IIIB (a) or BAL (b,c) or NL4-3 viral supernatants and BSA-Alexa 647 for 2 h at 37°C in the presence of CD4-Ig or sCD4 (a), the indicated anti-Env CD4-binding-site antibodies (b), or the indicated LCK inhibitors (c). Representative flow cytometry histograms of probe uptake in CD4+ T cells are shown at left. Graphs on the right show percent probe uptake in the presence of inhibitors relative to the no-inhibitor control from independent donors. Statistical significance was determined using a Student’s 1-sample t-test.

Env binding to CD4 could induce macropinocytosis directly by initiating CD4-mediated intracellular signaling that converges on iPLA2 and on pathways regulating actin polymerization, leading to macropinocytic cup formation. Alternatively, or in addition, Env conformational changes that occur on CD4 binding could expose regions of Env that permit Env binding to distinct receptors that mediate macropinocytosis. Concerning the former possibility, the CD4 cytoplasmic tail associates tightly with the LCK Src-family protein tyrosine kinase, which facilitates signaling through the TCR complex^47,48^. Therefore, we asked if Env-induced macropinocytosis could be blocked by a generic Src-family PTK inhibitor, PP2^49^, and a highly-specific LCK inhibitor, RK24466^47,50^. PP2 and RK24466 partially inhibited Env-induced macropinocytosis in CD4+ T cells, consistent with the idea that Env-induced macropinocytosis is in part mediated by CD4 signaling to T cells (Fig. 4c).

### Macropinocytosis induction in CD4+ T cells is a property of HIV-1 clade B Envs

We examined if soluble Envs from other HIV-1 clades could induce macropinocytosis in resting CD4+ T cells. All additional tested HIV-1 gp120 Envs of clade B viruses (B.MN, B.9021, B.63521, AC1029) induced probe uptake in resting CD4+ T cells, albeit to varying degrees (Fig. 5a). In contrast, gp120 Envs from clade AE viruses (CM235, 93TH975, AE.244) and clade C viruses (CN54, 96ZM651) were inactive (Fig. 5a). Thus, the ability of HIV-1 Env to induce CD4+ T cell macropinocytosis may be specific to Env from clade B viruses that are the predominant circulating strains in Europe, North and South America, and Australasia^51,52^. Nonetheless, an HIV-1 main (M) consensus gp120 Env (M.CONS), in which the most common amino acid across all HIV-1 M clades is represented at each position of the gp120 protein, was able to induce CD4+ T cell macropinocytosis (Fig. 5a). In addition, Env from simian immunodeficiency virus (SIV) was active (Fig. 5a).

**Figure 5.**
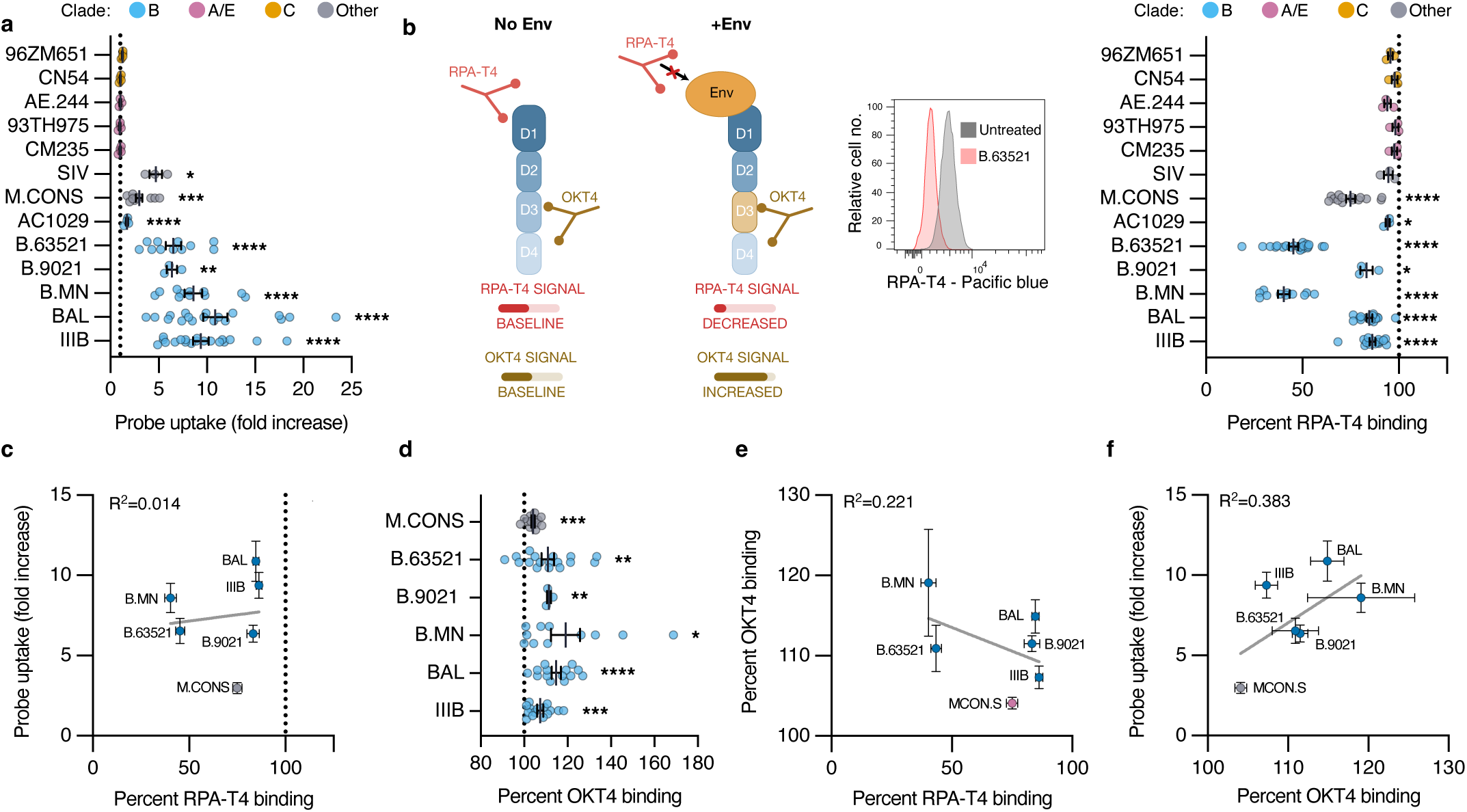
Env-induced macropinocytosis induction in CD4+ T cells is a property of HIV-1 clade B Envs and correlates with OKT4 binding to CD4 D3. **(a)** PBMC were incubated with soluble gp120 Envs from HIV-1 clade B (IIIB, BAL, B.MN, B.9021, B.63521, AC1029; blue), clade AE (CM235, 93TH975, AE.244; pink), clade C (CN54, 96ZM651; orange), an HIV-1 main (M) group consensus gp120 (M.CONS; gray), or SIV-1 Env (SIV; gray) (all 2 µg/ml), together with BSA-Alexa 647 for 2 h at 37°C. Shown is fold induction of BSA uptake in CD4+ T cells from independent donors. **(b-f)** PBMC were preincubated with soluble Envs (2 µg/ml) at 4°C, washed, and stained with the anti-CD4 D1 antibody, RPA-T4, and the anti-CD4 D3 antibody, OKT4. (**b**) A schematic of the effect of Env interaction with CD4 upon anti-CD4 antibody binding is shown at left. A representative flow cytometry histogram of the influence of B.63521 upon RPA-T4 binding to CD4+ T cells is shown in the middle panel. The graph at right shows the percent RPA-T4 binding after treatment with the indicated Env relative to the control without Env from independent donors. **(c)** Percent RPA-T4 binding plotted against fold induction of probe uptake in CD4+ T cells for the indicated Envs. **(d)** The graph shows the percent OKT4 binding after treatment with different Envs relative to the control without Env from independent donors. Statistical significance was determined using a Student’s 1-sample t-test (a,b,d). **(e)** Percent RPA-T4 binding versus OKT4 binding in CD4+ T cells for the indicated Envs. **(f)** Percent OKT4 binding versus fold induction of probe uptake in CD4+ T cells. Regression lines are shown with coefficients of determination (c,e,f).

Given the role of CD4 in macropinocytosis induction, we next asked if the different ability of Envs to induce macropinocytosis could be accounted for by differences in the extent to which they engage CD4 on the T cell surface. To assess this, T cells were preincubated with soluble Envs at 4°C, washed, and stained with the RPA-T4 antibody whose epitope on D1 of CD4 overlaps with the Env-binding epitope^6,53,54^. The extent of Env inhibition of RPA-T4 binding to CD4 was then determined by flow cytometry and taken as a measure of the strength of Env binding to CD4 (Fig. 5b). Significantly, soluble Envs from clade AE and clade C viruses, which did not induce macropinocytosis (Fig. 5a), only weakly inhibited RPA-T4 binding (Fig. 5b), indicating relatively low affinity for CD4. In contrast, Envs from all clade B viruses and SIV were generally more effective inhibitors of RPA-T4 binding, indicating stronger Env binding to CD4 (Fig. 5b). However, among the clade B Envs, the extent of CD4 binding did not correlate with the amount of macropinocytosis induction (Fig. 5c). For example, B.63521 and B.MN Envs showed much stronger interaction with CD4 than IIIB and BAL Envs, but were less potent inducers of macropinocytosis (Fig. 5c). Therefore, although Env interaction with CD4 is required for macropinocytosis, the degree of Env-CD4 binding does not necessarily predict its magnitude.

In all RPA-T4 binding-inhibition experiments conducted at 4°C, T cells were also stained with the OKT4 anti-CD4 antibody. The OKT4 epitope is located in D3 of CD4^55^, distinct from the RPA-T4 epitope, and was initially included as a control (Fig. 5b). However, in nearly all experiments conducted with clade B Envs, the decrease in RPA-T4 binding to D1 was accompanied by an increase in OKT4 binding to D3 (Fig. 5d and Supplementary Fig. 3). Plots of the mean percentage RPA-T4 binding versus the mean percentage OKT4 binding for different clade B Envs, did not reveal a clear correlation (Fig. 5e). In contrast, a stronger correlation was observed between the mean percentage OKT4 binding and the mean fold increase in probe uptake for clade B Envs (Fig. 5f). Notably, RPA-T4 itself did not induce macropinocytosis in CD4+ T cells at 37°C and did not influence the extent of binding of OKT4 to CD4 at 4°C (Supplementary Fig. 4). These findings suggest a model in which Env-specific induced conformational changes in CD4, which increase exposure of the OKT4 epitope, play an important role in the induction of macropinocytosis.

### Antibodies against the Env variable 3 loop and the bridging sheet inhibit Env-induced macropinocytosis

Upon Env interaction with CD4, Env conformational changes occur that expose the V3 loop of Env and result in the formation of the Env bridging sheet^5,7,8,56^. The V3 loop and bridging sheet subsequently bind to the CXCR4 or CCR5 chemokine receptors on target cells, leading to exposure of the gp41 Env fusion peptide and the fusion of viral and host membranes. To examine if the V3 loop is involved in the induction of CD4+ T cell macropinocytosis, we examined the effect of anti-V3 loop antibodies on BSA probe uptake. We focused upon IIIB, BAL and B.63521 and selected anti-V3 loop antibodies that are known to bind the V3 loops of these Envs (IIIB, 902; BAL, 447-52D, and 268 D IV; B.63521, 447-52D)^57–61^. All tested V3 antibodies inhibited Env-induced macropinocytosis (Fig. 6a). However, for IIIB and BAL, V3 antibodies also inhibited Env interaction with CD4, and correspondingly blocked Env-induced increased accessibility of D3, as determined in RPA-T4 and OKT4 binding assays conducted at 4°C (Fig. 6b-d). The B.63521 Env was an exception, as the 447-52D anti-V3 loop antibody partially inhibited probe uptake without affecting Env binding to CD4 (Fig. 6a-c). Nonetheless, 447-52D inhibited B.63521-induced OKT4 epitope accessibility (Fig. 6d). Altogether, these results are consistent with a model in which anti-V3 loop antibodies inhibit macropinocytosis, not by blocking interaction of the V3 loop with an unknown cell surface receptor, but by interfering with Env-induced CD4 signaling events.

**Figure 6.**
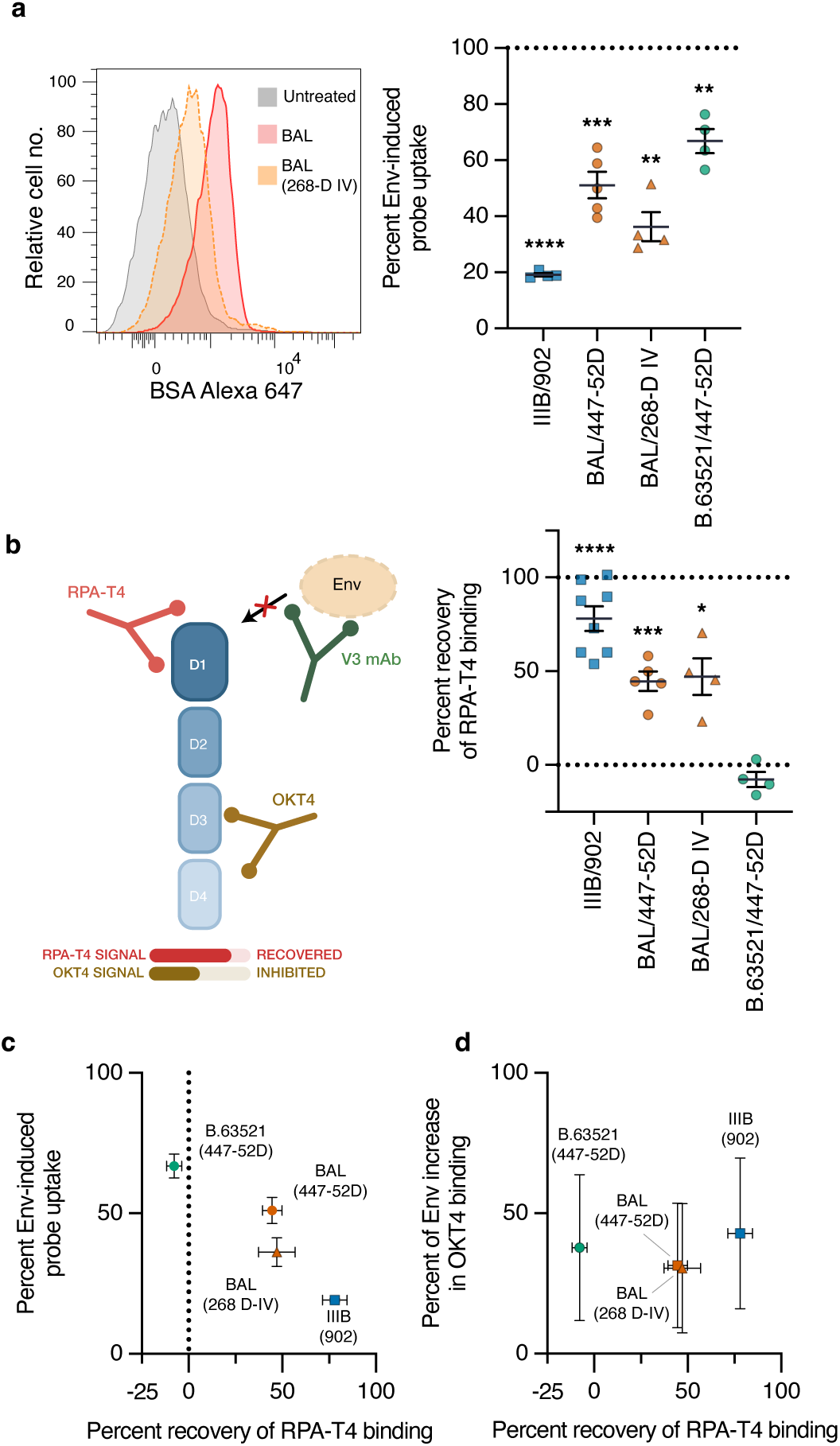
Effect of anti-V3 loop antibodies on Env-induced macropinocytosis, Env-CD4 binding, and Env-induced induction of OKT4 binding. **(a)** PBMC were incubated with IIIB, BAL, or B.63521 Env (2 µg/ml) in the presence of 902, 447-52D or 268-D IV anti-V3 loop antibodies together with BSA-Alexa 647 for 2 h at 37°C. A representative histogram showing BAL-induced probe uptake by CD4+ T cells in the presence and absence of 268-D IV is shown at left. The graph at right shows the percent Env-induced probe uptake in the presence of antibodies, relative to the absence of antibodies, across independent donors. **(b)** PBMC were incubated with Envs (2 µg/ml) in the presence or absence of anti-V3 loop antibodies at 4°C, washed, and stained with the anti-CD4 D1 antibody, RPA-T4, and the anti-CD4 D3 antibody, OKT4. The schematic at left shows the potential effect of anti-V3 loop antibodies on Env inhibition of RPA-T4 binding and increased OKT4 binding. The graph at right shows the percent anti-V3 loop antibody recovery of RPA-T4 binding in CD4+ T cells relative to Env in the absence of anti-V3 loop antibody (baseline) and CD4+ T cells without Env (100 percent recovery) from independent donors. Statistical significance was determined using a Student’s 1-sample t-test (a,b). **(c)** Plot of recovery of RPA-T4 binding versus percent Env-induced probe uptake for the indicated Envs and anti-V3 loop antibodies. (**d**) Plot of percent recovery of RPA-T4 binding versus percent increase of Env-induced OKT4 binding in the presence of anti-V3 loop antibodies relative to their absence.

We also examined the effect of antibodies against the Env bridging sheet on Env-induced CD4+ T cell macropinocytosis. Bridging sheet antibodies E51 and m9 (a single chain Fv antibody derived from the 17b antibody)^62–64^ inhibited macropinocytosis induced by IIIB, BAL, and B.63521 (Fig. 7a). Both antibodies partially inhibited IIIB binding to CD4 (Fig. 7b,c). However, the bridging sheet antibodies had only minor influences on the binding of BAL and B.63521 to CD4 (Fig. 7b,c). Nonetheless, E51 and m9 had an inhibitory influence upon the ability of BAL and B.63521 to induce the OKT4 epitope on CD4 (Fig. 7d). As with anti-V3 antibodies, these findings suggest that the inhibitory effect of anti-bridging sheet antibodies upon macropinocytosis is explained by their inhibition of Env-induced CD4 signaling rather than blockade of the interaction of Env with a distinct macropinocytosis-inducing cell surface receptor.

**Figure 7.**
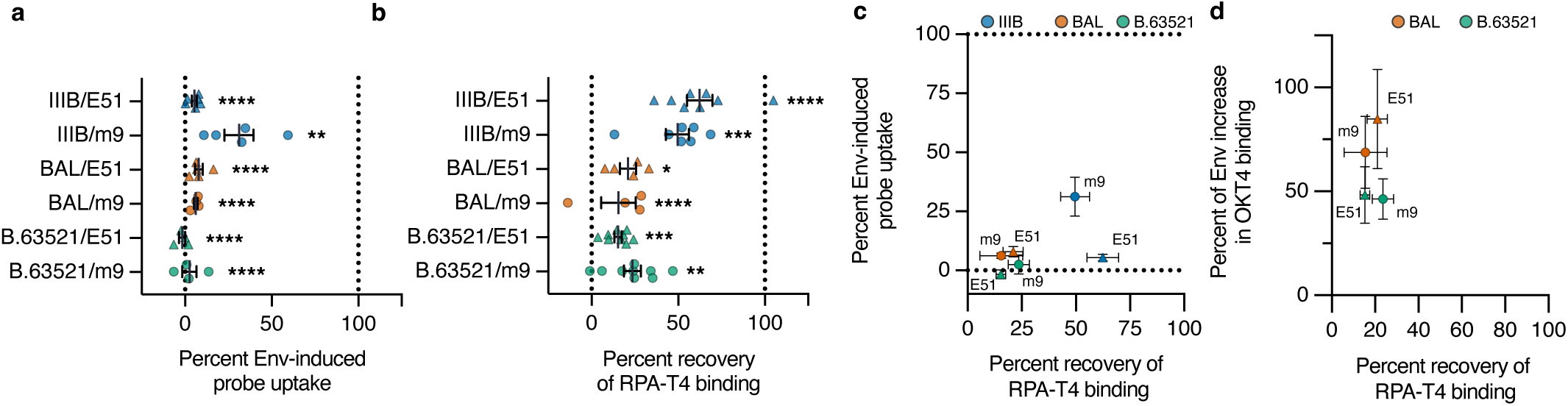
Effect of bridging sheet antibodies on Env-induced macropinocytosis, Env-CD4 binding, and Env-induced induction of OKT4 binding. **(a)** PBMC were incubated with IIIB, BAL, or B.63521 Env (2 µg/ml) in the presence of E51 or m9 anti-bridging sheet antibodies and BSA-Alexa 647 for 2 h at 37°C. Shown is the percent Env-induced probe uptake in the presence of antibodies, relative to the absence of antibodies, across independent donors. (**b**) PBMC were incubated with Envs (2 µg/ml) in the presence or absence of anti-bridging sheet antibodies at 4°C, washed, and stained with the anti-CD4 D1 antibody, RPA-T4, and the anti-CD4 D3 antibody, OKT4. Shown is the percent anti-bridging sheet antibody recovery of RPA-T4 binding in CD4+ T cells relative to Env in the absence of antibody (baseline) and CD4+ T cells without Env (100 percent recovery) from independent donors. Statistical significance was determined using a Student’s 1-sample t-test (a,b). **(c)** Plot of recovery of RPA-T4 binding versus percent Env-induced probe uptake for the indicated Envs and anti-bridging sheet antibodies. (**d**) Plot of percent recovery of RPA-T4 binding versus percent increase of Env-induced OKT4 binding in the presence of anti-bridging sheet antibodies relative to their absence.

### CXCR4 and other reported HIV-1 Env coreceptors are not required for Env-induced macropinocytosis in CD4+ T cells

We directly asked whether Env binding to chemokine coreceptors on resting T cells is necessary for Env-induced macropinocytosis. Of the two classical Env chemokine coreceptors, CXCR4 and CCR5, only CXCR4 is a relevant candidate, since CCR5 is expressed on a small minority of resting CD4+ T cells^65^, yet Env proteins induce essentially all resting CD4+ T cells to internalize BSA. To examine the potential role of CXCR4, we used the AMD3100 inhibitor that occupies the CRS2 pocket of CXCR4, thereby blocking V3 loop and Env interaction with the receptor^66–68^. At concentrations known to inhibit HIV infection of T cells, AMD3100 did not inhibit either soluble Env-induced or NL4-3-induced CD4+ T cell macropinocytosis (Supplementary Fig. 5a). Furthermore, an antibody against CXCR4 that blocks interaction with Env did not affect Env-induced macropinocytosis (Supplementary Fig. 5b).

Additional evidence against a role for CXCR4 included the findings that its natural ligand, CXCL12, neither induced CD4+ T cell macropinocytosis nor blocked Env-induced macropinocytosis of CD4+ T cells (Supplementary Fig. 5c,d). The only other chemokine receptor expressed on the majority of resting CD4+ T cells that could potentially engage the Env V3 loop and bridging sheet is CCR7^69^. However, an antibody against CCR7 did not affect Env-induced macropinocytosis (Supplementary Fig. 5e).

The V3 loop of Env has also been reported to bind glycosphingolipids (GSL), including lactosyl ceramide (LacCer) and the gangliosides GM3 and GD3^70–72^, each of which is present on the outer leaflet of the plasma membrane of resting human CD4+ T cells^73–75^. V3 loop interaction with gangliosides is potentially relevant since GM1 and Gb3 gangliosides have been shown to induce membrane curvature and invagination following their engagement by Cholera toxin B (GM1) SV40 (GM1) or Shiga toxin (Gb3)^76–78^. A required step in the synthesis of LacCer and all gangliosides except GM4 involves the addition of glucose to ceramide by the enzyme, GlcCer synthase^79,80^. Therefore, to examine a potential role for LacCer and gangliosides in Env-induced macropinocytosis, we tested the effect of the GlcCer synthase inhibitors, dl-threo-1-phenyl-2-(palmitoylamino)-3-morpholino-1-propanol (PPMP) and Genz-123346^81,82^. Both inhibitors potently blocked Env-induced macropinocytosis in CD4+ T cells (Supplementary Fig. 6a).

However, antibodies against LacCer, GM3 and GD3 had no effect on Env-induced macropinocytosis (Supplementary Fig. 6b). In addition, Cholera toxin B did not affect probe uptake (Supplementary Fig. 6c). GSL, along with cholesterol, are highly enriched in membrane lipid rafts, where a substantial fraction of CD4 resides^83–85^. Potentially, therefore, a role for GSL in Env-induced macropinocytosis reflects a role in lipid raft assembly rather than a role as a signaling molecule that binds Env.

### Env-induced macropinocytosis promotes HIV infection of CD4+ T cells

We have previously shown that macropinosomes are an important site of HIV-1 entry into the cytosol of activated CD4+ T cells, and that this route of entry contributes to HIV-1 infection of this cell type^30^. Therefore, the ability of HIV-1 Env to induce macropinocytosis in CD4+ T cells is likely to contribute to infection. To address this directly, we examined the effect of macropinocytosis inhibitors on the infection of resting CD4+ T cells. We previously showed that EIPA can inhibit HIV-1 infection of activated CD4+ T cells^30^. However, since EIPA blocks constitutive as well as Env-induced macropinocytosis in activated CD4+ T cells, this result does not specifically demonstrate a function for Env-induced macropinocytosis in infection. In contrast, essentially all macropinocytosis in resting human CD4+ T cells exposed to HIV-1 is Env-driven. Therefore, the effects of macropinocytosis inhibitors on the infection of resting T cells can be more straightforwardly interpreted. To assess infection, resting CD4+ T cells were incubated for 2 h with the NL4-3 virus encoding NanoLuc luciferase in the absence and presence of EIPA (Fig. 3a) and the PLA-2 inhibitors AACOCF3 and FKGK18 (Fig. 3b). Cells were then washed and incubated for a further 24 h before lysis and assessment of NanoLuc activity by chemiluminescence. In an initial experiment performed with four independent donors, we also examined the ability of NL4-3/NanoLuc to induce CD4+ T cell macropinocytosis. Across four independent donors, the extent of NL4-3/NanoLuc-induced macropinocytosis strongly correlated with the extent of NL4-3/NanoLuc infection, consistent with a role for Env-induced macropinocytosis in promoting infection (Fig. 8a). In support of this, each of EIPA, AACOCF3, and FKGK18 inhibited infection of resting CD4+ T cells (Fig. 8b).

**Figure 8.**
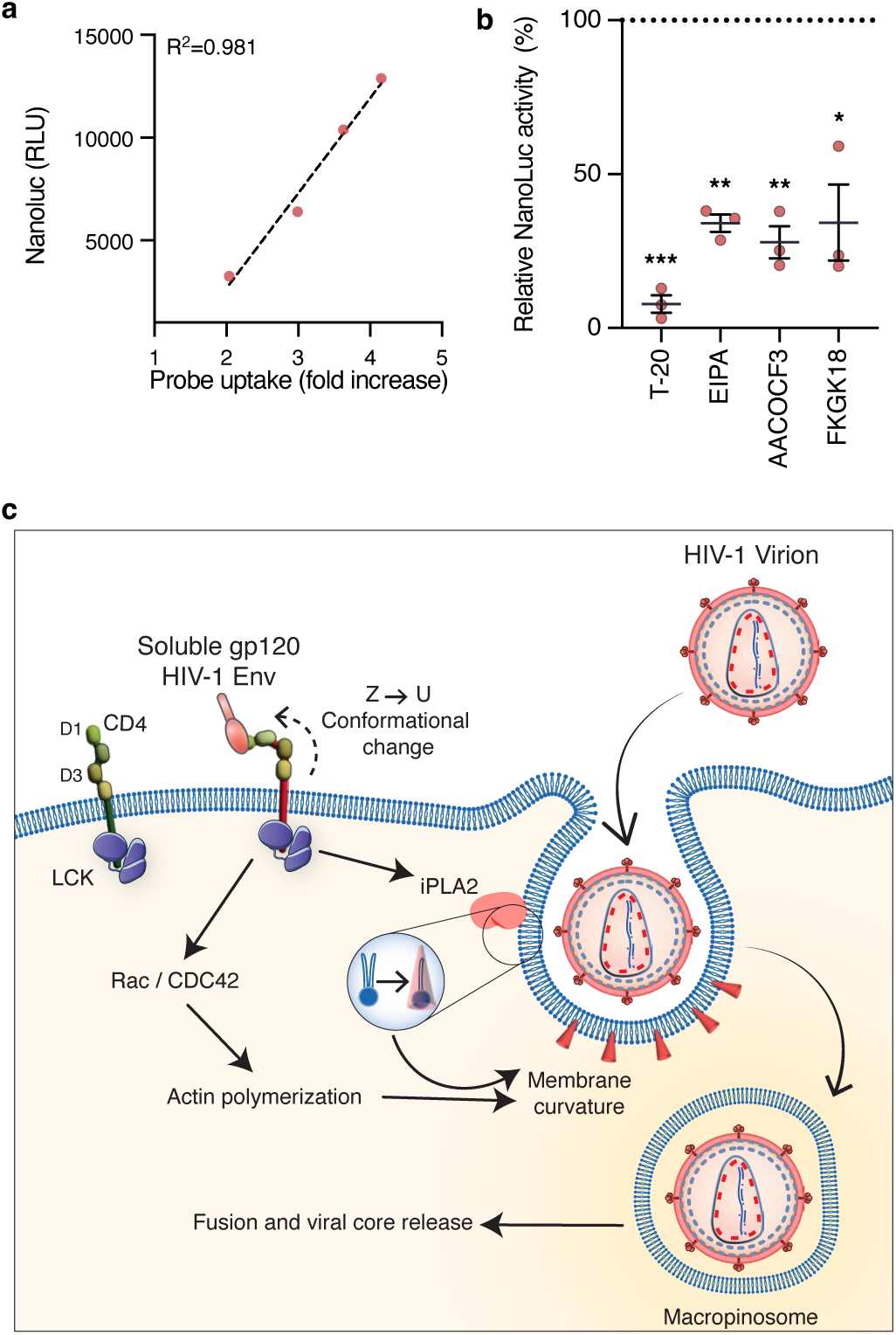
Env-induced macropinocytosis promotes HIV-1 infection of resting CD4+ T cells. **(a)** Resting CD4+ T cells from four different donors were inoculated with NL4-3/NanoLuc in the presence of BSA-Alexa 647 for 2 h at 37°C, before washing and determination of probe uptake by flow cytometry. Separate samples of resting CD4+ T cells from the same donors were incubated with NL4-3/NanoLuc without probe for 2 h at 37°C. Cells were then washed and incubated for a further 24 h in the presence of Saquinavir before lysis and assessment of NanoLuc activity by luminescence. Shown are fold increases in probe uptake versus NanoLuc counts for each donor. The regression line with the coefficient of determination is shown. **(b)** Resting CD4+ T cells were pre-incubated with the fusion inhibitor, T-20, or macropinocytosis inhibitors EIPA, AACOCF3, and FKGK18 for 15 min and incubated with NL4-3/NanoLuc for 2 h at 37°C in the continued presence of macropinocytosis inhibitors and Saquinavir, washed, and cultured for 24 h before lysis and determination of NL4-3/NanoLuc activity. The graph shows NanoLuc activity in the presence of inhibitor relative to the absence of inhibitor from independent donors. Statistical significance was determined using a Student’s 1-sample t-test. **(c)** Model of mechanism and role of HIV-1-induced macropinocytosis in resting CD4+ T cells. The binding of soluble HIV-1 gp120 Env to the CD4 ectodomain triggers CD4 conformational changes that activate iPLA-2, which converts phospholipids to cone-shaped lysophospholipids on the inner leaflet of the plasma membrane, resulting in inward membrane curvature. Additional signals from CD4 converge upon Rac/Cdc42 and the actin cytoskeleton, which promote further membrane curvature leading to macropinocytosis. Intact HIV-1 virions enter macropinosomes, where HIV-1 fusion with the membrane may be favored, resulting in productive infection.

## Discussion

We previously showed that macropinosomes are a significant route of HIV-1 entry into the cytoplasm of activated primary human CD4+ T cells, contributing to productive HIV-1 infection^30^. This property of HIV-1 is thus shared with several other viruses that are able to use macropinosomes as entry routes for the infection of their respective target cell types^23–27^. In the current studies, we sought to extend these findings by asking whether HIV-1 can itself induce macropinocytosis in CD4+ T cells, rather than passively being taken up by ongoing macropinocytosis that occurs as activated T cells progress through the cell cycle. HIV-1 was shown to trigger macropinocytosis in resting CD4+ T cells, leading to viral uptake into macropinosomes (Fig. 8c). As far as we are aware, the ability of HIV-1 to induce macropinocytosis in CD4+ T cells is unique among viruses that use macropinocytosis as a cell entry route.

Further analyses indicated that soluble gp120 Env, likely shed from HIV-1 virions, and not Env tethered to the HIV-1 virion, is responsible for macropinocytosis induction. This was supported by the finding that virions purified from crude supernatants had much reduced activity, whereas post-spin supernatants retained activity, and by the finding that crude supernatants from Env-negative viruses also lacked activity. Furthermore, soluble recombinant gp120 Env was a potent inducer of macropinocytosis in resting CD4+ T cells and augmented macropinocytosis in already activated CD4+ T cells. Activity was largely confined to clade B Envs, which is nonetheless significant given that clade B viruses are the dominant strains that are actively circulating in Europe, the Americas, and Australasia, especially in specific risk groups^51,52^. It is unclear why virion-tethered Env was less able to induce macropinocytosis in our experiments, but this is unlikely to be a consequence of its trimeric nature since soluble trimeric Env was shown to be an active inducer of CD4+ T cell macropinocytosis. It is possible that the lack of activity is a consequence of an artefactual detachment of Env from the virion surface during the process of sucrose centrifugation, resulting in low amounts of Env. Alternatively, the number of trimers present on the virion surface, estimated at 7-12^86–88^, may be insufficient for macropinocytosis. Regardless, the soluble gp120 Env concentrations used in the current studies are physiologically relevant. Concentrations of soluble gp120 in the serum of untreated HIV-1-infected patients have been estimated at approximately 0.1 to 1 µg/ml^36,37^, within the range of concentrations of soluble gp120 shown to induce CD4+ T cell macropinocytosis in this study.

To address the biological significance of Env-induced macropinocytosis, we examined the influence of inhibitors of macropinocytosis upon HIV-1 infection of resting CD4+ T cells. Resting CD4+ T cells were chosen for this purpose since they engage in minimal macropinocytosis prior to incubation with Env. Therefore, any inhibitory effect of macropinocytosis inhibitors on the infection of resting CD4+ T cells can be interpreted as indicating that Env-induced macropinocytosis specifically contributes to infection. Significantly, EIPA, an established specific inhibitor of macropinocytosis, and AACOCF3 and FKGK18, two novel inhibitors of macropinocytosis described herein, both of which target PLA-2, inhibited HIV-1 infection of resting CD4+ T cells. These findings thus illustrate the significance of Env-induced macropinocytosis for infection, at least in resting CD4+ T cells (Fig. 8c).

It is interesting to speculate why HIV-1 has evolved to induce macropinocytosis in resting CD4+ T cells to facilitate HIV-1 infection of this cell type. In our previous studies, we demonstrated that macropinocytosis in T cells delivers key extracellular amino acids to lysosomes, that is necessary for the activation of mTORC at the lysosomal surface^28^. This is potentially significant, as mTORC is known to promote productive HIV-1 infection of CD4+ T cells through multiple mechanisms^89–91^. These include stimulation of the pentose phosphate shunt pathway, which increases the availability of dNTPs required for viral reverse transcription; inhibition of viral autophagy; and activation of lipogenesis, which facilitates viral assembly and budding. Full activation of mTORC, as occurs in activated T cells progressing through the cell cycle, requires two signals. The first signal is provided by the TCR and CD28, which trigger the phosphatidylinositol 3-kinase(PI3K)/protein kinase B (PKB)/tuberous sclerosis (TSC) pathway that culminates in the activation of Rheb GTPases. The second signal is provided by TCR/CD28-induced macropinocytic delivery of extracellular amino acids to lysosomes, which activates Rag GTPases^28,29^. In contrast to TCR/CD28, HIV-1 Env induces a transient activation of mTORC in resting CD4+ T cells^92,93^. It has been shown previously that HIV-1 can activate the PI3K/PKB/TSC pathway in resting CD4+ T cells, thus providing the first signal for mTORC activation^92–94^. Our discovery here that HIV-1 Env triggers macropinocytosis in CD4+ T cells constitutes the second signaling arm that could account for the transient mTORC activation observed previously. We propose that this transient mTORC activation is an important factor that explains why certain HIV strains have evolved to induce macropinocytosis in resting CD4+ T cells to facilitate their own infection.

Aside from mTORC activation, we showed recently that HIV-1 can fuse with the macropinosome membrane and gain access to the cytoplasm from this site in activated CD4+ T cells^30^. Similarly, others had shown previously that HIV-1 fusion in CD4+ T cells occurs predominantly within endosomes^10,12^. Therefore, another possible explanation for Env-induced macropinocytosis is that it permits access of HIV-1 to an environment that is more permissive for fusion than the plasma membrane. Following their excision from the plasma membrane, macropinosomes undergo shrinkage and tubulation, which permit the recycling of endocytosed membrane to the cell surface^95,96^. During shrinkage, the membrane density of CD4 and chemokine coreceptors would be expected to increase relative to their plasma membrane levels. In turn, this increased density of receptor and coreceptor could promote fusion and HIV-1 entry.

As evidenced by the blocking effect of soluble CD4 D1-containing proteins and antibodies against the Env CD4-binding site, Env-induced macropinocytosis is dependent upon Env-CD4 interaction. In addition, antibodies against the Env V3 loop and bridging sheet inhibited Env-induced macropinocytosis, initially suggesting that macropinocytosis might also depend upon Env interaction with HIV-1 plasma membrane coreceptors. However, we found that anti-V3 and bridging sheet antibodies frequently destabilized Env interaction with CD4, possibly as a result of steric hindrance, but nonetheless providing an alternative explanation for macropinocytosis inhibition. Furthermore, we found no evidence of a requirement for known HIV-1 coreceptors in Env-induced macropinocytosis. Therefore, we propose that Env interaction with CD4 is sufficient to induce macropinocytosis in CD4+ T cells (Fig. 8c). As shown by small-angle X-ray scattering, Env binding induces structural changes in the CD4 extracellular region characterized by a transition from a semi-extended tetralobal Z conformation to a U bi-fold conformation, likely owing to Env-induced alterations in the flexible D2-D3 linker of the CD4^97^. These changes are unique to Env interaction with CD4 and are not observed with tick saliva protein that also binds CD4 D1^97^. The unique structural changes could trigger activation of downstream signaling events that couple CD4 to macropinocytosis. In this study, the notion that Env induces conformational changes in CD4 on the surface of CD4+ T cells is supported by our observation that Env incubation at 4°C results in increased binding of the OKT4 antibody that binds D3 of CD4. Increased OKT4 binding is consistent with a Z-to-U transition of CD4 that is expected to increase the accessibility OKT4 to D3. Moreover, evidence that CD4 conformational changes are integral to the induction of macropinocytosis is provided by our finding that the ability of different clade B Envs to increase OKT4 binding correlates with their ability to induce macropinocytosis. This is in contrast to their ability to inhibit the binding of the RPA-T4 CD4 antibody, a measure of the strength of Env-CD4 interaction, that does not correlate with the extent of macropinocytosis. Furthermore, the RPA-T4 antibody itself does not increase OKT4 binding to CD4+ T cells and, correspondingly, does not induce macropinocytosis, despite the fact that it binds an epitope on D1 that partially overlaps with the Env interaction epitope. Last, for the B.63521 Env, the 447-52D anti-V3 loop antibody inhibited macropinocytosis but did not affect CD4 binding, and for the B.63521 and BAL Envs, the E51 and m9 anti-bridging sheet antibodies inhibited macropinocytosis but had only a small inhibitory effect on CD4 binding. However, in each of these instances, the antibody inhibited the ability of Env to increase OKT4 binding. This inhibition could potentially be explained by the ability of antibodies to restrain further conformational changes in Env, necessary for the CD4 Z-to-U transition. However, regardless of the mechanism, the fact that the antibodies blocked both Env-induced macropinocytosis and the induction of the OKT4 epitope, provides further evidence of a role for CD4 conformational changes in the triggering of this event.

An intriguing finding in the current study is that PLA-2 is required for Env-induced macropinocytosis in resting CD4+ T cells as well as all other forms of macropinocytosis examined (Fig. 8c). PLA-2s catalyze the hydrolysis of membrane phospholipids at their sn-2 position to generate free fatty acids and lysophospholipids^98^. PLA-2s comprise a large family of enzymes with at least 16 different subfamilies^98^. However, in CD4+ T cells, expression of non-secreted cytosolic forms of PLA-2 is largely restricted to group 6 calcium-independent iPLA-2 forms, specifically group 6A (beta, PNPLA9), 6B (gamma, PNPLA8), 6C (delta, PNPLA6), 6E (zeta, PNPLA2) and 6F (eta, PNPLA4 isoform)^99^. Thus, any or all of these isoforms could participate in Env-induced macropinocytosis, although it is noted that FKGK11 and FKGK18 inhibitors exhibit high specificity for iPLA-2 beta^44,45^. PLA-2 has not been implicated in macropinocytosis previously. However, PLA-2 has been shown to be necessary for human monocyte phagocytosis of antibody-coated erythrocytes and for mitogen-driven human lymphocyte proliferation^100–102^. In the former case, blocked phagocytosis following PLA-2 inhibition could be rescued by the addition of arachidonic acid. In contrast, we have been unable to rescue Env-induced macropinocytosis with arachidonic acid. Therefore, we propose that it is the generation of lysophospholipids by PLA-2 rather than fatty acids that promotes Env-induced macropinocytosis. As a result of their inverted cone-like structure, lysophospholipids can induce membrane curvature when asymmetrically distributed in the lipid bilayer in the direction they are most concentrated^103,104^. Significantly, in early studies, HIV-1 Env was shown to activate PLA-2 in human monocytes and T cells upon interaction with CD4^105,106^. This activated PLA-2 could thus act on phospholipids in the inner leaflet of the plasma membrane to generate lysophospholipids at this site, which could initiate membrane curvature and macropinocytosis.

Interestingly, the crystal structure of iPLA-2 beta has revealed a dimeric molecule in which ankyrin repeat domains contribute to closure of the active site and distance the catalytic domains from membrane substrates^107^. Activation of iPLA-2 beta by displacement of bound calmodulin is proposed to open the catalytic site and position it closer to the plasma membrane. The initial generation of lysophospholipid would then cause membrane curvature, which could facilitate further interaction of enzyme and substrate, resulting in further membrane bending in a positive feedback loop that leads to full macropinocytosis. However, it is important to point out that although necessary, PLA-2 activation is not sufficient for Env-induced macropinocytosis since antibodies against CD4 D1 have also been shown to induce PLA-2 activation^105,106^, yet do not induce macropinocytosis. Given the demonstrated role of Env-induced macropinocytosis in HIV-1 infection of CD4+ T cells, how Env-induced CD4 conformational changes, LCK, PLA-2, and Rho family GTPases coordinate to orchestrate this process should be the focus of future investigation.

## Methods

### Human samples, ethics, and biosafety

Human leukocyte packs from healthy adult donors were obtained from the New York Blood Center. Products were CPD-anticoagulated leukocyte preparations and were processed for PBMC isolation within 24 h of collection. All donors were deidentified and no donor-specific inclusion or exclusion criteria beyond the healthy-donor source were applied. Unless otherwise stated, each data point shown in the study represents one unique donor. All work involving HIV-1 molecular clones, virus-containing supernatants, and primary-cell infection assays was conducted under BSL-2+ containment in accordance with institutional biosafety procedures.

### Cells and culture conditions

293T cells (ATCC, cat. CRL-3216) were used for transient transfection and production of HIV-1-containing supernatants. Cells were maintained at 37°C in 5% CO₂ in D10 medium: DMEM (Gibco, cat. 11995073) supplemented with 10% FBS (Gibco, cat. 26140079), 100 U/mL penicillin, 100 µg/mL streptomycin (Invitrogen, cat. 15140122), and 1× GlutaMAX (Gibco, cat. 35050061). PBMC, purified CD4+ T cells, and purified peripheral blood T cell preparations were cultured in R10 medium: RPMI 1640 (Gibco, cat. 11875093) supplemented with 10% FBS, 100 U/mL penicillin, 100 µg/mL streptomycin, and 1× GlutaMAX. PHA-activated T cells were maintained in R10 supplemented with 50 IU/mL recombinant human IL-2 (BD Biosciences, cat. 554603).

### Isolation of PBMC and primary T-cell populations

PBMC were isolated from NYBC leukocyte products by density-gradient centrifugation using Ficoll-Paque PLUS (Cytiva, cat. 17144002) in SepMate-50 tubes (Stemcell Tech, cat. 85460) according to manufacturer’s instructions. Purified resting CD4+ T cells were isolated from PBMC by magnetic negative selection using the Miltenyi Human CD4+ T Cell Isolation Kit (130-096-533) according to the manufacturer’s instructions. Where pan-T-cell preparations were required, the Miltenyi Human Pan T Cell Isolation Kit (130-096-535) was used. Where activated T cells were examined, cells were stimulated with phytohemagglutinin (PHA-P, Invitrogen, 6µg/mL) for approximately 24 h, washed, and then used in macropinocytosis or assays.

### HIV-1 molecular clones, recombinant Env proteins, and anti-Env reagents

The HIV-1 molecular clones used in this study included NL4-3, NL4-3 lacking Env (ΔEnv), and an NL4-3 NanoLuc reporter virus. These constructs were used in HIV-1 entry and infection studies as previously^30^.

Recombinant HIV-1 Env proteins and anti-Env reagents were obtained through the NIH HIV Reagent Program. Env proteins used in the present study included gp120 IIIB (ARP-11784), gp120 BaL (HRP-20082), gp140 trimer SF162 (ARP-12026), B.MN Δ11 gp120 (ARP-12570), B.9021 Δ11 gp120 (ARP-12571), B.63521 Δ11 gp120 mutC (ARP-12574), AE.A244 Δ11 gp120 (ARP-12569), AC10.29 gp120 (ARP-13055), 96ZM651 gp120 (ARP-10080), CM235 gp120 (ARP-12816), 93TH975 gp120 (ARP-13342), CN54 gp120 (ARP-13354), M.CON-S Δ11 gp120 (ARP-12576), and SIVmac239 gp130-His (ARP-12797). Proteins were stored at −80°C in the supplied buffer and thawed once for use. Unless otherwise indicated, soluble Env proteins were used at 2 μg/mL.

Anti-Env reagents included CD4-Ig (ARP-13058, Human CD4-Ig Recombinant Protein), soluble CD4 (sCD4; ARP-4615), and CD4 binding site antibodies, N6 (ARP-12968), NIH45-46 G54W (ARP-12174), and 654-30D (ARP-7369). Anti-Env V3-loop antibodies used included 902 (ARP-522), 447-52D (ARP-4030), and 268-D IV (ARP-1511). Bridging-sheet antibodies included E51 (ARP-11439) and m9 (ARP-11710). V3-loop and bridging-sheet antibodies were used at 1 μg/mL.

### Additional ligands, antibodies, and inhibitors

Anti-CXCR4 antibody (1µg/mL; R&D Systems, cat. MAB172), CXCL12 (50 ng/mL; Gibco, cat. AF-300-28A-10), anti-CCR7 antibody (1µg/mL; R&D Systems, cat. MAB197), cholera toxin B subunit (1µg/mL; Invitrogen, cat. C34775), anti-GM3 antibody (2 µg/mL; Novus Biologicals, cat. NB469372), anti-GD3 antibodies (1µg/mL; MilliporeSigma, clone UM4D4 cat. MABF978 and clone R24 cat. MABC1112), anti-GalCer antibody (1 µg/mL; MilliporeSigma, Cat. No. MAB342), anti-LacCer antibody (1µg/mL; Novus Biologicals, clone HULY-m13, cat. NBP2-45291), and anti-GM1 antibody (1µg/mL; Sigma-Aldrich, cat. SAE0069) were used as indicated.

Inhibitors used in the study included EIPA (50µM; Sigma-Aldrich, cat. A3085), jasplakinolide (1µM; Cayman Chemical, cat. 11705), blebbistatin (75µM; Cayman Chemical, cat. 13013), EHT1864 (20µM; Selleck Chemical, cat. S7482), ML141 (10µM; Selleck Chemical, Cat. S7686), PitStop2 (25µM; Cayman Chemical, cat. 23885), AACOCF3 (50µM; Cayman Chemical, cat. 62120), BEL (100µM; Cayman Chemical, cat. 70700), FKGK11 (88µM; Cayman Chemical, cat. 13179), FKGK18 (100µM; Cayman Chemical, cat. 13943), pyrrophenone (2µM; Cayman Chemical, cat. 13294), PP2 (10µM; Selleck Chemical, Cat. S7008), RK24466 (10µM; MedChemExpress, cat. HY-108318), AMD3100 (10µM; Selleck Chemical, cat. S3013), PPMP (20µM; Cayman Chemical, cat. 22677), Genz-123346 (25µM; Cayman Chemical, cat. 28500), and T-20 (2µM; NIH HIV Reagent Program). Stocks were prepared in DMSO or the vendor-recommended solvent, and matched vehicle controls were included in each experiment.

### Virus production, clarification, ultracentrifugation, and normalization

Virus-containing supernatants were generated by transient transfection of 293T cells with HIV-1 molecular clones using Lipofectamine 2000 (Invitrogen). Briefly, 5 × 10⁶ 293T cells were plated on 10 cm tissue culture dishes and cultured for 1 day at 37°C prior to transfection with 20 µg of the relevant molecular clone. Two days post-transfection, supernatants were clarified by filtration through a 0.45 µm filter and either used directly in uptake assays, frozen as single-use aliquots at −80°C, or subjected to ultracentrifugation through sucrose.

For virion-purification experiments, clarified virus-containing supernatants were layered over a 20% sucrose cushion and ultracentrifuged at 24,000 rpm for 2 h at 4°C in a TH641 rotor (ThermoFisher). After centrifugation, the post-spin supernatant was carefully collected above the sucrose cushion, avoiding disturbance of the virion pellet, and retained as the virus-depleted supernatant. Virion pellets were resuspended in RPMI containing 10% FBS, aliquoted, and stored at −80°C. Virus-depleted supernatants were aliquoted and stored at −80°C until use.

Virus input was normalized by p24 amount, which was quantified by ELISA as described previously^30^. Briefly, viral particles were lysed in ELISA lysis buffer (0.05% Tween 20, 0.5% Triton X-100, 0.5% casein in PBS). An anti-HIV-1 p24 antibody (clone 183-H12-5C; NIH HIV Reagent Program) was bound to Nunc MaxiSorp plates overnight at 4°C. Lysed samples were incubated on plates for 2 h, and captured p24 was detected by sequential incubation with a biotinylated anti-HIV-1 p24 antibody (clone 31-90-25; ATCC), streptavidin-HRP (Fitzgerald), and TMB substrate (Sigma). Biotinylation was performed using the EZ-Link Micro Sulfo-NHS-Biotinylation Kit (Pierce). p24 concentrations were determined against recombinant HIV-1 IIIB p24 protein standards (NIH HIV Reagent Program). In all macropinocytosis and infection assays, an amount of virus that contained 5 µg of p24 was used.

### Flow cytometry

Flow cytometric detection of macropinocytosis was performed as described previously^28^. PBMC or purified T-cell preparations (2–3 × 10^6^ cells) were incubated in 500 μL RPMI with 80µg/mL fluorescent BSA (BSA-Alexa Fluor 647 ThermoFisher cat. A34785; BSA-Alexa Fluor 488 ThermoFisher cat. A13100) and the indicated stimuli for 2 h at 37°C in FACS tubes (Falcon, 14-959-2A). Inhibitors and antibodies were added to cultures 30 min prior to the addition of the probe. Following washes, cells were stained with OKT4-Alexa Fluor 488 (BioLegend 317420) and PE/Cyanine7 anti-human CD8 (BioLegend 344712). Samples were acquired on BD FACSCanto and/or BD LSRFortessa flow cytometers and analyzed using FlowJo v10 or v11. The standard gating sequence for lymphocyte analyses proceeded from FSC/SSC to the lymphocyte gate, followed by singlet discrimination and then separation of CD4+ and CD8+ T-cell populations. Monocytes within PBMC were identified using forward light scatter and CD4 staining. Fold increase of probe uptake was calculated as the ratio of the median fluorescence intensity (MFI) of stimulated versus unstimulated cells. Percent Env-induced uptake in the presence of inhibitors was calculated as [MFI (Env plus inhibitor) – MFI (unstimulated)/MFI (Env alone) – MFI (unstimulated)] × 100.

To detect the influence of Env on the binding of anti-CD4 antibodies to CD4, PBMC were incubated or not with soluble Env proteins at 2 μg/mL at 4°C for 30 min, washed, and stained with CD4 antibodies, RPA-T4-Pacific Blue (BioLegend 300521) and/or OKT4-AF488. CD4 staining of gated CD4+ T cells was then analyzed by flow cytometry. All staining and wash steps in these assays were performed at 4°C. Percent RPA-T4 binding was calculated as [MFI (Env treatment) / MFI (without Env treatment] × 100. In antibody-recovery experiments (Figs. 6–7), anti-Env antibodies (1 μg/mL) were included in the Env preincubation step. Percent antibody recovery of RPA-T4 binding was calculated as [MFI (Env plus anti-Env antibody) − MFI (Env treatment) / MFI (without Env treatment) − MFI (Env treatment)] × 100. Percent OKT4 binding was calculated as [MFI (Env treatment) / MFI (without Env treatment] × 100. The effect of anti-Env antibodies on OKT4 antibody binding was expressed as the percent of Env increase in OKT4 binding and was calculated as [MFI (Env plus anti-Env antibody) - MFI (without Env treatment) / MFI (Env treatment) - MFI (without Env treatment)] × 100.

### Microscopy of BSA-positive vesicles and virion localization

For imaging assays, purified resting T cells or CD4+ T cells were first subjected to the BSA probe uptake workflow described above. Cells were washed 3x in ice-cold PBS (300 x g, 4 min, 4°C) and then allowed to settle onto 0.1% poly-L-lysine-coated (Millipore Sigma, S4521) 12mm #1.5 round glass coverslips (Thomas Scientific 1217N79) for 30 mins at 4°C. Cells were fixed with 4% paraformaldehyde for 20 min at room temperature, permeabilized and blocked using PBS containing 0.1% saponin, 3% BSA, and 5% goat serum for 30 min at room temperature. To detect HIV-1 particles within BSA-positive vesicles, cells were stained with an anti-HIV-1 p24 capsid protein antibody (AG3.0, NIH HIV Reagent Program, ARP-4121) followed by goat anti-mouse IgG1-Alex 568 F(ab) secondary antibody (Invitrogen, A66783; 1:400). Samples were additionally stained with OKT4-Alexa Fluor 488 (BioLegend 317420) or Alexa Fluor 488 anti-human CD8 (BioLegend 344716). Coverslips were mounted in ProLong Diamond anti-fade (Invitrogen, P36971) and images were acquired by spinning-disk confocal microscopy (Nikon CSU-X1) with a 100× objective. Z-stacks were collected at 200nm intervals where needed to visualize vesicular localization. Acquisition settings were kept constant between different samples within an experiment. Representative images were processed using Nikon NIS-Elements software, with downstream image processing in ImageJ/Fiji.

### NanoLuc infection assays

Resting purified CD4+ T cells (1 × 10⁶) were plated in 96-well U-bottom plates and pretreated with inhibitors or vehicle controls for 30 min at 37°C. Cells were then inoculated with 5µg of p24 NL4-3 NanoLuc reporter viruses for 2 h at 37°C. Inhibitors were maintained during the inoculation period. Cells were washed to remove unbound virus and inhibitors, and cultured for an additional 24 h in the presence of 10µM Saquinavir (NIH HIV Reagent program) to restrict replication to a single cycle, before lysis with 1× Passive Lysis Buffer (PLB; Promega). NanoLuc activity in lysates was measured using the Nano-Glo Luciferase Assay System (Promega) and normalized to total protein as determined by the Pierce 660 nm Protein Assay Reagent (Pierce). Relative NanoLuc activity in the presence of inhibitors was expressed as a percentage of vehicle-treated control.

### Statistical analysis

Data are presented as mean ± SEM. Statistical analyses and graphing were performed in GraphPad Prism 11. The Student’s 1-sample *t*-test was used for comparisons to theoretical means, the Mann-Whitney *U* test for pairwise comparisons, and the Kruskal-Wallis test with Dunn’s correction for multiple comparisons. Linear regression was used to assess the correlation between parameters in data sets. The following symbols were used to denote statistical significance: * *p*<0.05, ** *p*<0.01, *** *p*<0.001, **** *p*<0.0001.

## Acknowledgements

This work was supported by NIH grant R01 AI65381 to P.D.K. and A.O.

## Author contributions

P.M., A.O., and P.D.K. designed experiments that were carried out by P.M., C.SdC., Y.T., X.W., T.M, and P.D.K. P.M. and P.D.K. wrote the manuscript with input from all authors.

---

**Supplementary Figure 1.** Trimeric gp140 Env induces CD4+ T cell macropinocytosis. PBMC were incubated with BSA-Alexa 488 and soluble trimeric gp140 from HIV-1 strain SF162 (clade B, CCR5 tropic) or monomeric gp120 IIIB for 2 h at 37°C, or with SF162 at 4°C as a negative control (all 2 µg/ml or at concentrations otherwise indicated). Representative flow cytometry histograms showing BSA uptake in CD4+ and CD8+ T cells are shown. The graph at right shows fold induction of SF162-induced BSA probe uptake from independent donors (2 µg/ml of SF162). Statistical significance was determined using a Mann-Whitney *U* test.

**Supplementary Figure 2.** Inhibition of monocyte and activated T cell macropinocytosis by inhibitors of calcium-independent PLA-2. **(a)** PBMC were incubated with BSA-Alexa 647 for 2 h at 37°C in the presence of the PLA-2 inhibitors, arachidonyl trifluoromethyl ketone (AACOCF3), bromoenol lactone (BEL), FKGK11, FKGK18, or pyrrophenone (Pyro). Representative flow cytometry histograms of probe uptake by CD4+ monocytes are shown at left. Graphs on the right show the percent probe uptake in the presence of inhibitors compared to the no-inhibitor control from independent donors. **(b)** PHA-activated PBMC were incubated with BSA-Alexa 647 for 2 h at 37°C in the presence of AACOCF3 or FKGK18. Representative flow cytometry histograms of probe uptake by CD4+ and CD8+ T cells are shown at top. Graphs at bottom show the percent probe uptake in the presence of inhibitors compared to the no-inhibitor control from independent donors. Statistical significance was determined using a Student’s 1-sample t-test.

**Supplementary Figure 3.** Effect of soluble gp120 Envs upon binding of anti-CD4 D1 and D3 antibodies. PBMC were incubated with the indicated Envs (2 µg/ml) at 4°C, washed, and stained with the anti-CD4 D1 antibody, RPA-T4, and the anti-CD4 D3 antibody, OKT4. Data points in plots show the percent RPA-T4 binding versus percent OKT4 binding on CD4+ T cells in the presence of Env relative to the absence of Env from different donors.

**Supplementary Figure 4.** The RPA-T4 antibody does not induce CD4+ T cell macropinocytosis and does not increase the binding of the OKT4 antibody to CD4. **(a)** PBMC were incubated with the anti-CD4 D1 antibody RPA-T4 or gp120 IIIB for 2 h at 37°C in the presence of BSA-Alexa 647. Shown is the fold induction of BSA probe uptake in CD4+ T cells from independent donors. **(b)** PBMC were incubated with RPA-T4 at 4° C, washed, and stained with OKT4. Percent OKT4 binding relative to untreated CD4+ T cells is shown. Statistical significance was determined using a Mann-Whitney *U* test.

**Supplementary Figure 5.** Chemokine receptor inhibitors, antibodies, and agonists do not inhibit Env-induced macropinocytosis in CD4+ T cells. (a-e) PBMC were incubated with soluble gp120 IIIB or BAL Envs (2 µg/ml) or NL4-3 virus-containing supernatants in the presence of BSA-Alexa 647 for 2 h at 37°C in the presence of AMD3100 (CXCR4 inhibitor) (a), an anti-CXCR4 antibody that blocks Env interaction with CXCR4 (b), CXCL12 (SDF-1; CXCR4 ligand) (c,d), or an anti-CCR7 antibody that blocks CCL19 and CCL21 binding (e). Graphs show the percent probe uptake by CD4+ T cells in the presence of inhibitors, antibodies, and chemokines compared to the no-inhibitor control from independent donors (a,b,d,e) or the fold increase in CD4+ T cell probe uptake induced by CXCL12 or BAL from independent donors (c). Statistical significance was determined using a Student’s 1-sample t-test.

**Supplementary Figure 6.** Effect of inhibitors of GSL synthesis and anti-GSL reagents on Env-induced macropinocytosis in CD4+ T cells. **(a)** PBMC were pretreated with GlcCer synthase inhibitors PPMP or Genz-123346 for 24 hours, before incubation with gp120 IIIB or BAL (2 µg/ml) and BSA-Alexa 647 for 2 h 37°C. Percent Env-induced BSA probe uptake in inhibitor-treated CD4+ T cells relative to CD4+ T cells without inhibitor pretreatment is shown for independent donors. **(b)** PBMC were incubated with BAL (2 µg/ml) and BSA-Alexa 647 for 2 h at 37°C in the presence of antibodies against GAL-Cer, LacCer and gangliosides, GM3 and GD3. Shown is the percent BAL-induced probe uptake in CD4+ T cells in the presence of antibodies compared to CD4+ T cells in the absence of antibodies from independent donors. **(c)** PBMC were incubated with BAL (2 µg/ml) and BSA-Alexa 647 for 2 h at 37°C in the presence of cholera toxin B subunit (GM1-binding). Shown is the percent BAL-induced probe uptake in CD4+ T cells in the presence of CTB compared to CD4+ T cells in the absence of CTB from independent donors. Statistical significance was determined using a Student’s 1-sample t-test.

## References

1 Deeks, S. G., Overbaugh, J., Phillips, A. & Buchbinder, S. HIV infection. Nat Rev Dis Primers 1, 15035 (2015). 10.1038/nrdp.2015.35

2 Maartens, G., Celum, C. & Lewin, S. R. HIV infection: epidemiology, pathogenesis, treatment, and prevention. Lancet 384, 258–271 (2014). 10.1016/S0140-6736(14)60164-1

3 Chen, B. Molecular Mechanism of HIV-1 Entry. Trends Microbiol 27, 878–891 (2019). 10.1016/j.tim.2019.06.002

4 Wilen, C. B., Tilton, J. C. & Doms, R. W. HIV: cell binding and entry. Cold Spring Harb Perspect Med 2 (2012). 10.1101/cshperspect.a006866

5 Dam, K. A., Fan, C., Yang, Z. & Bjorkman, P. J. Intermediate conformations of CD4-bound HIV-1 Env heterotrimers. Nature 623, 1017–1025 (2023). 10.1038/s41586-023-06639-8

6 Kwong, P. D. et al. Structure of an HIV gp120 envelope glycoprotein in complex with the CD4 receptor and a neutralizing human antibody. Nature 393, 648–659 (1998). 10.1038/31405

7 Li, W. et al. HIV-1 Env trimers asymmetrically engage CD4 receptors in membranes. Nature 623, 1026–1033 (2023). 10.1038/s41586-023-06762-6

8 Shaik, M. M. et al. Structural basis of coreceptor recognition by HIV-1 envelope spike. Nature 565, 318–323 (2019). 10.1038/s41586-018-0804-9

9 Carter, G. C., Bernstone, L., Baskaran, D. & James, W. HIV-1 infects macrophages by exploiting an endocytic route dependent on dynamin, Rac1 and Pak1. Virology 409, 234–250 (2011). 10.1016/j.virol.2010.10.018

10 de la Vega, M. et al. Inhibition of HIV-1 endocytosis allows lipid mixing at the plasma membrane, but not complete fusion. Retrovirology 8, 99 (2011). 10.1186/1742-4690-8-99

11 Marechal, V. et al. Human immunodeficiency virus type 1 entry into macrophages mediated by macropinocytosis. J Virol 75, 11166–11177 (2001). 10.1128/JVI.75.22.11166-11177.2001

12 Sharma, M., Marin, M., Wu, H., Prikryl, D. & Melikyan, G. B. Human Immunodeficiency Virus 1 Preferentially Fuses with pH-Neutral Endocytic Vesicles in Cell Lines and Human Primary CD4+ T-Cells. ACS Nano 17, 17436–17450 (2023). 10.1021/acsnano.3c05508

13 Yasen, A., Herrera, R., Rosbe, K., Lien, K. & Tugizov, S. M. HIV internalization into oral and genital epithelial cells by endocytosis and macropinocytosis leads to viral sequestration in the vesicles. Virology 515, 92–107 (2018). 10.1016/j.virol.2017.12.012

14 Liu, N. Q. et al. Human immunodeficiency virus type 1 enters brain microvascular endothelia by macropinocytosis dependent on lipid rafts and the mitogen-activated protein kinase signaling pathway. J Virol 76, 6689–6700 (2002). 10.1128/jvi.76.13.6689-6700.2002

15 Herold, N. et al. HIV-1 entry in SupT1-R5, CEM-ss, and primary CD4+ T cells occurs at the plasma membrane and does not require endocytosis. J Virol 88, 13956–13970 (2014). 10.1128/JVI.01543-14

16 Bloomfield, G. & Kay, R. R. Uses and abuses of macropinocytosis. J Cell Sci 129, 2697–2705 (2016). 10.1242/jcs.176149

17 Buckley, C. M. & King, J. S. Drinking problems: mechanisms of macropinosome formation and maturation. FEBS J 284, 3778–3790 (2017). 10.1111/febs.14115

18 Kerr, M. C. & Teasdale, R. D. Defining macropinocytosis. Traffic 10, 364–371 (2009). 10.1111/j.1600-0854.2009.00878.x

19 Liu, Z. & Roche, P. A. Macropinocytosis in phagocytes: regulation of MHC class-II-restricted antigen presentation in dendritic cells. Front Physiol 6, 1 (2015). 10.3389/fphys.2015.00001

20 Schuette, V. & Burgdorf, S. The ins-and-outs of endosomal antigens for cross-presentation. Curr Opin Immunol 26, 63–68 (2014). 10.1016/j.coi.2013.11.001

21 Commisso, C. et al. Macropinocytosis of protein is an amino acid supply route in Ras-transformed cells. Nature 497, 633–637 (2013). 10.1038/nature12138

22 Palm, W. et al. The Utilization of Extracellular Proteins as Nutrients Is Suppressed by mTORC1. Cell 162, 259–270 (2015). 10.1016/j.cell.2015.06.017

23 de Vries, E. et al. Dissection of the influenza A virus endocytic routes reveals macropinocytosis as an alternative entry pathway. PLoS Pathog 7, e1001329 (2011). 10.1371/journal.ppat.1001329

24 Mercer, J. & Helenius, A. Vaccinia virus uses macropinocytosis and apoptotic mimicry to enter host cells. Science 320, 531–535 (2008). 10.1126/science.1155164

25 Mercer, J. & Helenius, A. Virus entry by macropinocytosis. Nat Cell Biol 11, 510–520 (2009). 10.1038/ncb0509-510

26 Nanbo, A. et al. Ebolavirus is internalized into host cells via macropinocytosis in a viral glycoprotein-dependent manner. PLoS Pathog 6, e1001121 (2010). 10.1371/journal.ppat.1001121

27 Shema Mugisha, C., et al. A Simplified Quantitative Real-Time PCR Assay for Monitoring SARS-CoV-2 Growth in Cell Culture. mSphere 5 (2020). 10.1128/mSphere.00658-20

28 Charpentier, J. C. et al. Macropinocytosis drives T cell growth by sustaining the activation of mTORC1. Nat Commun 11, 180 (2020). 10.1038/s41467-019-13997-3

29 Charpentier, J. C. & King, P. D. Mechanisms and functions of endocytosis in T cells. Cell Commun Signal 19, 92 (2021). 10.1186/s12964-021-00766-3

30 Murakami, T. et al. Macropinosomes are a site of HIV-1 entry into primary CD4(+) T cells. Proc Natl Acad Sci U S A 122, e2417676122 (2025). 10.1073/pnas.2417676122

31 Koivusalo, M. et al. Amiloride inhibits macropinocytosis by lowering submembranous pH and preventing Rac1 and Cdc42 signaling. J Cell Biol 188, 547–563 (2010). 10.1083/jcb.200908086

32 Nofal, M., Zhang, K., Han, S. & Rabinowitz, J. D. mTOR Inhibition Restores Amino Acid Balance in Cells Dependent on Catabolism of Extracellular Protein. Mol Cell 67, 936–946 e935 (2017). 10.1016/j.molcel.2017.08.011

33 Rychert, J., Strick, D., Bazner, S., Robinson, J. & Rosenberg, E. Detection of HIV gp120 in plasma during early HIV infection is associated with increased proinflammatory and immunoregulatory cytokines. AIDS Res Hum Retroviruses 26, 1139–1145 (2010). 10.1089/aid.2009.0290

34 Santosuosso, M., Righi, E., Lindstrom, V., Leblanc, P. R. & Poznansky, M. C. HIV-1 envelope protein gp120 is present at high concentrations in secondary lymphoid organs of individuals with chronic HIV-1 infection. J Infect Dis 200, 1050–1053 (2009). 10.1086/605695

35 Wyatt, R. et al. The antigenic structure of the HIV gp120 envelope glycoprotein. Nature 393, 705–711 (1998). 10.1038/31514

36 Cummins, N. W., Rizza, S. A. & Badley, A. D. How much gp120 is there? J Infect Dis 201, 1273–1274; author reply 1274-1275 (2010). 10.1086/651434

37 Oh, S. K. et al. Identification of HIV-1 envelope glycoprotein in the serum of AIDS and ARC patients. J Acquir Immune Defic Syndr (1988) 5, 251–256 (1992).

38 Merk, A. & Subramaniam, S. HIV-1 envelope glycoprotein structure. Curr Opin Struct Biol 23, 268–276 (2013). 10.1016/j.sbi.2013.03.007

39 Yoshida, S., Pacitto, R., Yao, Y., Inoki, K. & Swanson, J. A. Growth factor signaling to mTORC1 by amino acid-laden macropinosomes. J Cell Biol 211, 159–172 (2015). 10.1083/jcb.201504097

40 Shutes, A. et al. Specificity and mechanism of action of EHT 1864, a novel small molecule inhibitor of Rac family small GTPases. J Biol Chem 282, 35666–35678 (2007). 10.1074/jbc.M703571200

41. Surviladze, Z., et al. in Probe Reports from the NIH Molecular Libraries Program (2010).

42 von Kleist, L. et al. Role of the clathrin terminal domain in regulating coated pit dynamics revealed by small molecule inhibition. Cell 146, 471–484 (2011). 10.1016/j.cell.2011.06.025

43 Ackermann, E. J., Conde-Frieboes, K. & Dennis, E. A. Inhibition of macrophage Ca(2+)-independent phospholipase A2 by bromoenol lactone and trifluoromethyl ketones. J Biol Chem 270, 445–450 (1995). 10.1074/jbc.270.1.445

44 Kokotos, G. et al. Potent and selective fluoroketone inhibitors of group VIA calcium-independent phospholipase A2. J Med Chem 53, 3602–3610 (2010). 10.1021/jm901872v

45 Ali, T. et al. Characterization of FKGK18 as inhibitor of group VIA Ca2+-independent phospholipase A2 (iPLA2beta): candidate drug for preventing beta-cell apoptosis and diabetes. PLoS One 8, e71748 (2013). 10.1371/journal.pone.0071748

46 Ono, T. et al. Characterization of a novel inhibitor of cytosolic phospholipase A2alpha, pyrrophenone. Biochem J 363, 727–735 (2002). 10.1042/0264-6021:3630727

47 Gaud, G., Lesourne, R. & Love, P. E. Regulatory mechanisms in T cell receptor signalling. Nat Rev Immunol 18, 485–497 (2018). 10.1038/s41577-018-0020-8

48 Rudd, C. E. How the Discovery of the CD4/CD8-p56(lck) Complexes Changed Immunology and Immunotherapy. Front Cell Dev Biol 9, 626095 (2021). 10.3389/fcell.2021.626095

49 Hanke, J. H. et al. Discovery of a novel, potent, and Src family-selective tyrosine kinase inhibitor. Study of Lck- and FynT-dependent T cell activation. J Biol Chem 271, 695–701 (1996). 10.1074/jbc.271.2.695

50 Seo, H. H. et al. 7-cyclopentyl-5-(4-phenoxyphenyl)-7H-pyrrolo[2,3-d] pyrimidin-4-ylamine inhibits the proliferation and migration of vascular smooth muscle cells by suppressing ERK and Akt pathways. Eur J Pharmacol 798, 35–42 (2017). 10.1016/j.ejphar.2017.02.004

51 Bbosa, N., Kaleebu, P. & Ssemwanga, D. HIV subtype diversity worldwide. Curr Opin HIV AIDS 14, 153–160 (2019). 10.1097/COH.0000000000000534

52 Williams, A. et al. Geographic and Population Distributions of Human Immunodeficiency Virus (HIV)-1 and HIV-2 Circulating Subtypes: A Systematic Literature Review and Meta-analysis (2010-2021). J Infect Dis 228, 1583–1591 (2023). 10.1093/infdis/jiad327

53 Hong, Y. L. et al. New reporter cell lines to study macrophage-tropic HIV envelope protein-mediated cell-cell fusion. AIDS Res Hum Retroviruses 15, 1667–1672 (1999). 10.1089/088922299309702

54 Helling, B. et al. A specific CD4 epitope bound by tregalizumab mediates activation of regulatory T cells by a unique signaling pathway. Immunol Cell Biol 93, 396–405 (2015). 10.1038/icb.2014.102

55 Merkenschlager, M., Buck, D., Beverley, P. C. & Sattentau, Q. J. Functional epitope analysis of the human CD4 molecule. The MHC class II-dependent activation of resting T cells is inhibited by monoclonal antibodies to CD4 regardless whether or not they recognize epitopes involved in the binding of MHC class II or HIV gp120. J Immunol 145, 2839–2845 (1990).

56 Klasse, P. J., Sanders, R. W., Ward, A. B., Wilson, I. A. & Moore, J. P. The HIV-1 envelope glycoprotein: structure, function and interactions with neutralizing antibodies. Nat Rev Microbiol 23, 734–752 (2025). 10.1038/s41579-025-01206-6

57 Beaumont, T., Quakkelaar, E., van Nuenen, A., Pantophlet, R. & Schuitemaker, H. Increased sensitivity to CD4 binding site-directed neutralization following in vitro propagation on primary lymphocytes of a neutralization-resistant human immunodeficiency virus IIIB strain isolated from an accidentally infected laboratory worker. J Virol 78, 5651–5657 (2004). 10.1128/JVI.78.11.5651-5657.2004

58 Basu, S. et al. Determination of Binding Affinity of Antibodies to HIV-1 Recombinant Envelope Glycoproteins, Pseudoviruses, Infectious Molecular Clones, and Cell-Expressed Trimeric gp160 Using Microscale Thermophoresis. Cells 13 (2023). 10.3390/cells13010033

59 Conley, A. J. et al. Neutralization of primary human immunodeficiency virus type 1 isolates by the broadly reactive anti-V3 monoclonal antibody, 447-52D. J Virol 68, 6994–7000 (1994). 10.1128/JVI.68.11.6994-7000.1994

60 Killikelly, A. et al. Thermodynamic signatures of the antigen binding site of mAb 447-52D targeting the third variable region of HIV-1 gp120. Biochemistry 52, 6249–6257 (2013). 10.1021/bi400645e

61 Lusso, P. et al. Cryptic nature of a conserved, CD4-inducible V3 loop neutralization epitope in the native envelope glycoprotein oligomer of CCR5-restricted, but not CXCR4-using, primary human immunodeficiency virus type 1 strains. J Virol 79, 6957–6968 (2005). 10.1128/JVI.79.11.6957-6968.2005

62 Huang, C. C. et al. Structural basis of tyrosine sulfation and VH-gene usage in antibodies that recognize the HIV type 1 coreceptor-binding site on gp120. Proc Natl Acad Sci U S A 101, 2706–2711 (2004). 10.1073/pnas.0308527100

63 Zhang, M. Y. et al. Potent and broad neutralizing activity of a single chain antibody fragment against cell-free and cell-associated HIV-1. MAbs 2, 266–274 (2010). 10.4161/mabs.2.3.11416

64 Zhang, M. Y. et al. Improved breadth and potency of an HIV-1-neutralizing human single-chain antibody by random mutagenesis and sequential antigen panning. J Mol Biol 335, 209–219 (2004). 10.1016/j.jmb.2003.09.055

65 Lee, B., Sharron, M., Montaner, L. J., Weissman, D. & Doms, R. W. Quantification of CD4, CCR5, and CXCR4 levels on lymphocyte subsets, dendritic cells, and differentially conditioned monocyte-derived macrophages. Proc Natl Acad Sci U S A 96, 5215–5220 (1999). 10.1073/pnas.96.9.5215

66 Donzella, G. A. et al. AMD3100, a small molecule inhibitor of HIV-1 entry via the CXCR4 co-receptor. Nat Med 4, 72–77 (1998). 10.1038/nm0198-072

67 Sang, X. et al. Structural mechanisms underlying the modulation of CXCR4 by diverse small-molecule antagonists. Proc Natl Acad Sci U S A 122, e2425795122 (2025). 10.1073/pnas.2425795122

68 Zhang, Z. et al. CXCR4 mediated recognition of HIV envelope spike and inhibition by CXCL12. Nat Commun 16, 8653 (2025). 10.1038/s41467-025-63815-2

69 Campbell, J. J. et al. CCR7 expression and memory T cell diversity in humans. J Immunol 166, 877–884 (2001). 10.4049/jimmunol.166.2.877

70 Delezay, O., Hammache, D., Fantini, J. & Yahi, N. SPC3, a V3 loop-derived synthetic peptide inhibitor of HIV-1 infection, binds to cell surface glycosphingolipids. Biochemistry 35, 15663–15671 (1996). 10.1021/bi961205g

71 Hammache, D. et al. Specific interaction of HIV-1 and HIV-2 surface envelope glycoproteins with monolayers of galactosylceramide and ganglioside GM3. J Biol Chem 273, 7967–7971 (1998). 10.1074/jbc.273.14.7967

72 Planes, R. & Bahraoui, E. HIV and SIV Envelope Glycoproteins Interact with Glycolipids and Lipids. Int J Mol Sci 24 (2023). 10.3390/ijms241411730

73 Garofalo, T. et al. Association of GM3 with Zap-70 induced by T cell activation in plasma membrane microdomains: GM3 as a marker of microdomains in human lymphocytes. J Biol Chem 277, 11233–11238 (2002). 10.1074/jbc.M109601200

74 Sorice, M. et al. Evidence for the existence of ganglioside-enriched plasma membrane domains in human peripheral lymphocytes. J Lipid Res 38, 969–980 (1997).

75 Tani-ichi, S. et al. Structure and function of lipid rafts in human activated T cells. Int Immunol 17, 749–758 (2005). 10.1093/intimm/dxh257

76 Ewers, H. et al. GM1 structure determines SV40-induced membrane invagination and infection. Nat Cell Biol 12, 11–18; sup pp 11–12 (2010). 10.1038/ncb1999

77 Kabbani, A. M., Raghunathan, K., Lencer, W. I., Kenworthy, A. K. & Kelly, C. V. Structured clustering of the glycosphingolipid GM1 is required for membrane curvature induced by cholera toxin. Proc Natl Acad Sci U S A 117, 14978–14986 (2020). 10.1073/pnas.2001119117

78 Romer, W. et al. Shiga toxin induces tubular membrane invaginations for its uptake into cells. Nature 450, 670–675 (2007). 10.1038/nature05996

79 Groux-Degroote, S., Rodriguez-Walker, M., Dewald, J. H., Daniotti, J. L. & Delannoy, P. Gangliosides in Cancer Cell Signaling. Prog Mol Biol Transl Sci 156, 197–227 (2018). 10.1016/bs.pmbts.2017.10.003

80 Marks, D. L. et al. Oligomerization and topology of the Golgi membrane protein glucosylceramide synthase. J Biol Chem 274, 451–456 (1999). 10.1074/jbc.274.1.451

81 Abe, A. et al. Improved inhibitors of glucosylceramide synthase. J Biochem 111, 191–196 (1992). 10.1093/oxfordjournals.jbchem.a123736

82 Lee, L., Abe, A. & Shayman, J. A. Improved inhibitors of glucosylceramide synthase. J Biol Chem 274, 14662–14669 (1999). 10.1074/jbc.274.21.14662

83 Foti, M., Phelouzat, M. A., Holm, A., Rasmusson, B. J. & Carpentier, J. L. p56Lck anchors CD4 to distinct microdomains on microvilli. Proc Natl Acad Sci U S A 99, 2008–2013 (2002). 10.1073/pnas.042689099

84 Fragoso, R. et al. Lipid raft distribution of CD4 depends on its palmitoylation and association with Lck, and evidence for CD4-induced lipid raft aggregation as an additional mechanism to enhance CD3 signaling. J Immunol 170, 913–921 (2003). 10.4049/jimmunol.170.2.913

85 Lingwood, D. & Simons, K. Lipid rafts as a membrane-organizing principle. Science 327, 46–50 (2010). 10.1126/science.1174621

86 Chertova, E. et al. Envelope glycoprotein incorporation, not shedding of surface envelope glycoprotein (gp120/SU), Is the primary determinant of SU content of purified human immunodeficiency virus type 1 and simian immunodeficiency virus. J Virol 76, 5315–5325 (2002). 10.1128/jvi.76.11.5315-5325.2002

87 Zhu, P. et al. Electron tomography analysis of envelope glycoprotein trimers on HIV and simian immunodeficiency virus virions. Proc Natl Acad Sci U S A 100, 15812–15817 (2003). 10.1073/pnas.2634931100

88 Zhu, P. et al. Distribution and three-dimensional structure of AIDS virus envelope spikes. Nature 441, 847–852 (2006). 10.1038/nature04817

89 Crater, J. M., Nixon, D. F. & Furler O’Brien, R. L. HIV-1 replication and latency are balanced by mTOR-driven cell metabolism. Front Cell Infect Microbiol 12, 1068436 (2022). 10.3389/fcimb.2022.1068436

90 Taylor, H. E. et al. mTOR Overcomes Multiple Metabolic Restrictions to Enable HIV-1 Reverse Transcription and Intracellular Transport. Cell Rep 31, 107810 (2020). 10.1016/j.celrep.2020.107810

91 Heredia, A. et al. Targeting of mTOR catalytic site inhibits multiple steps of the HIV-1 lifecycle and suppresses HIV-1 viremia in humanized mice. Proc Natl Acad Sci U S A 112, 9412–9417 (2015). 10.1073/pnas.1511144112

92 Akbay, B., Shmakova, A., Vassetzky, Y. & Dokudovskaya, S. Modulation of mTORC1 Signaling Pathway by HIV-1. Cells 9 (2020). 10.3390/cells9051090

93 Wiredja, D. D. et al. Global phosphoproteomics of CCR5-tropic HIV-1 signaling reveals reprogramming of cellular protein production pathways and identifies p70-S6K1 and MK2 as HIV-responsive kinases required for optimal infection of CD4+ T cells. Retrovirology 15, 44 (2018). 10.1186/s12977-018-0423-4

94 Vasiliver-Shamis, G., Cho, M. W., Hioe, C. E. & Dustin, M. L. Human immunodeficiency virus type 1 envelope gp120-induced partial T-cell receptor signaling creates an F-actin-depleted zone in the virological synapse. J Virol 83, 11341–11355 (2009). 10.1128/JVI.01440-09

95 Freeman, S. A. et al. Lipid-gated monovalent ion fluxes regulate endocytic traffic and support immune surveillance. Science 367, 301–305 (2020). 10.1126/science.aaw9544

96 Zeziulia, M., Blin, S., Schmitt, F. W., Lehmann, M. & Jentsch, T. J. Proton-gated anion transport governs macropinosome shrinkage. Nat Cell Biol 24, 885–895 (2022). 10.1038/s41556-022-00912-0

97 Ashish et al. Conformational rearrangement within the soluble domains of the CD4 receptor is ligand-specific. J Biol Chem 283, 2761–2772 (2008). 10.1074/jbc.M708325200

98 Burke, J. E. & Dennis, E. A. Phospholipase A2 structure/function, mechanism, and signaling. J Lipid Res 50 **Suppl**, S237–242 (2009). 10.1194/jlr.R800033-JLR200

99 Uhlen, M. et al. A genome-wide transcriptomic analysis of protein-coding genes in human blood cells. Science 366 (2019). 10.1126/science.aax9198

100 Karimi, K., Gemmill, T. R. & Lennartz, M. R. Protein kinase C and a calcium-independent phospholipase are required for IgG-mediated phagocytosis by Mono-Mac-6 cells. J Leukoc Biol 65, 854–862 (1999). 10.1002/jlb.65.6.854

101 Lennartz, M. R. et al. Phospholipase A2 inhibition results in sequestration of plasma membrane into electronlucent vesicles during IgG-mediated phagocytosis. J Cell Sci 110 **(Pt** **17**), 2041–2052 (1997). 10.1242/jcs.110.17.2041

102 Roshak, A. K., Capper, E. A., Stevenson, C., Eichman, C. & Marshall, L. A. Human calcium-independent phospholipase A2 mediates lymphocyte proliferation. J Biol Chem 275, 35692–35698 (2000). 10.1074/jbc.M002273200

103 Fanani, M. L. & Ambroggio, E. E. Phospholipases and Membrane Curvature: What Is Happening at the Surface? Membranes (Basel) 13 (2023). 10.3390/membranes13020190

104 Fuller, N. & Rand, R. P. The influence of lysolipids on the spontaneous curvature and bending elasticity of phospholipid membranes. Biophys J 81, 243–254 (2001). 10.1016/S0006-3495(01)75695-0

105 Mavoungou, E. et al. HIV and SIV envelope glycoproteins induce phospholipase A2 activation in human and macaque lymphocytes. J Acquir Immune Defic Syndr Hum Retrovirol 16, 1–9 (1997). 10.1097/00042560-199709010-00001

106 Wahl, L. M. et al. Human immunodeficiency virus glycoprotein (gp120) induction of monocyte arachidonic acid metabolites and interleukin 1. Proc Natl Acad Sci U S A 86, 621–625 (1989). 10.1073/pnas.86.2.621

107 Malley, K. R. et al. The structure of iPLA(2)beta reveals dimeric active sites and suggests mechanisms of regulation and localization. Nat Commun 9, 765 (2018). 10.1038/s41467-018-03193-0

