## Supplementary figures and images for "The gp120 Envelope Glycoprotein of HIV-1 Triggers Macropinocytosis in Primary CD4+ T Cells to Promote HIV-1 Infection"

### Supplemental Figures

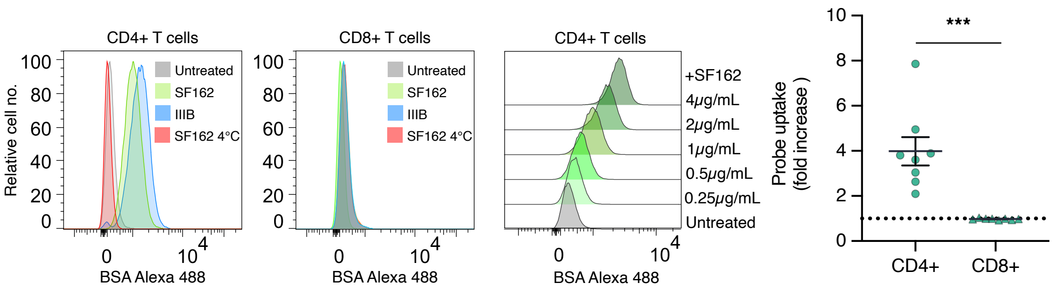

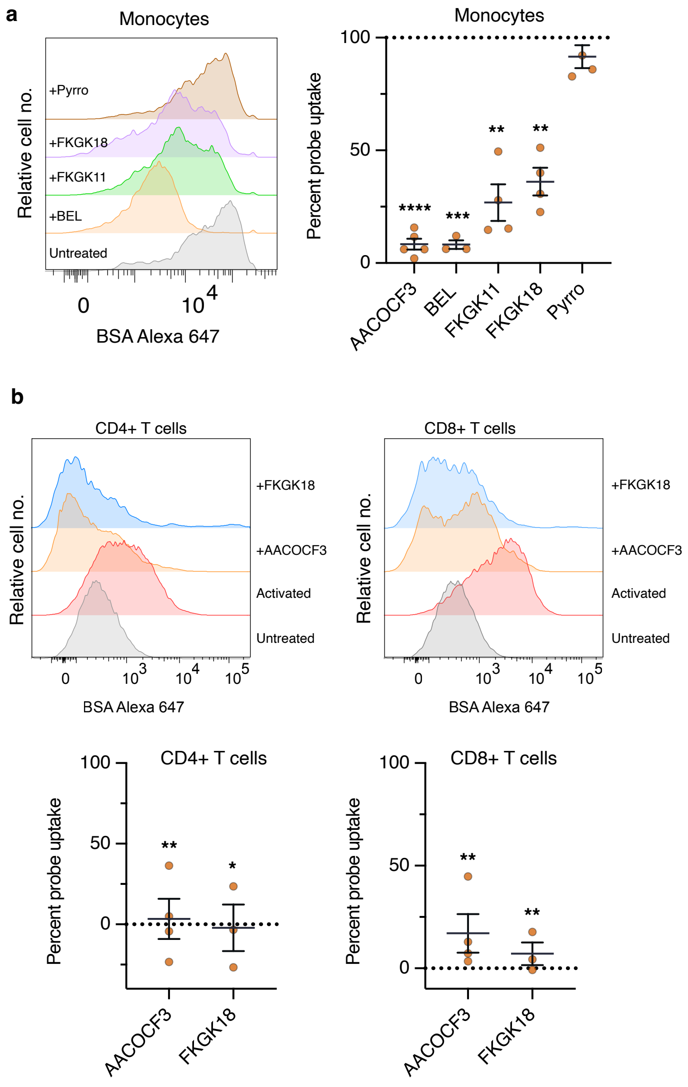

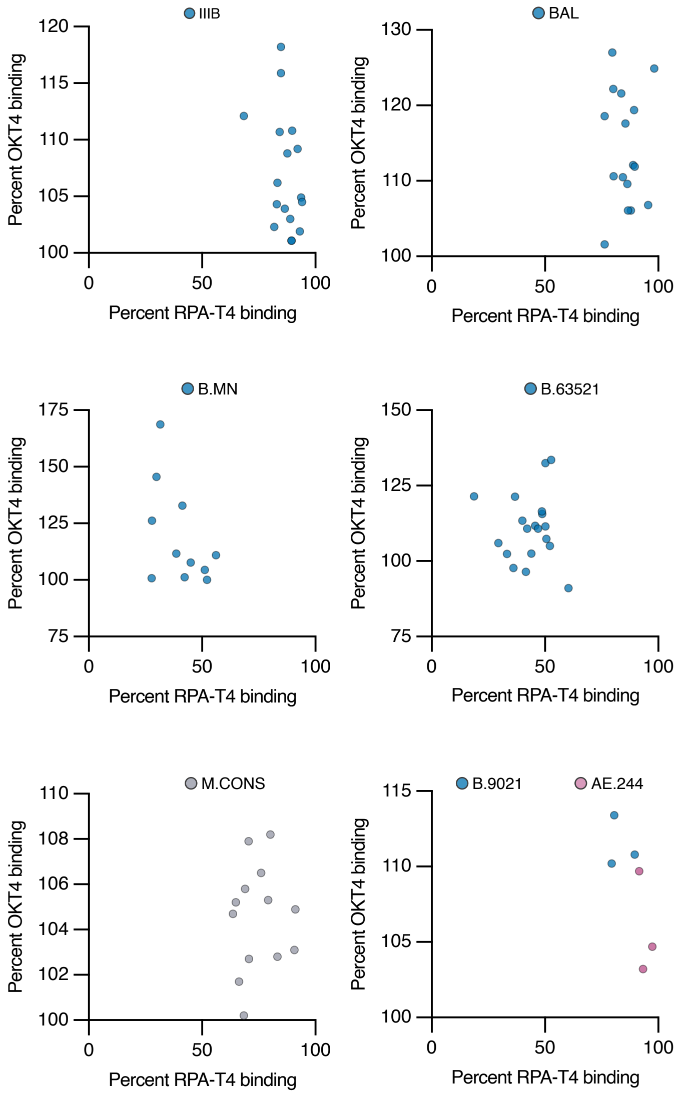

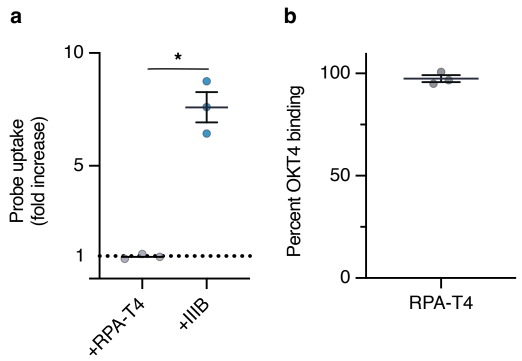

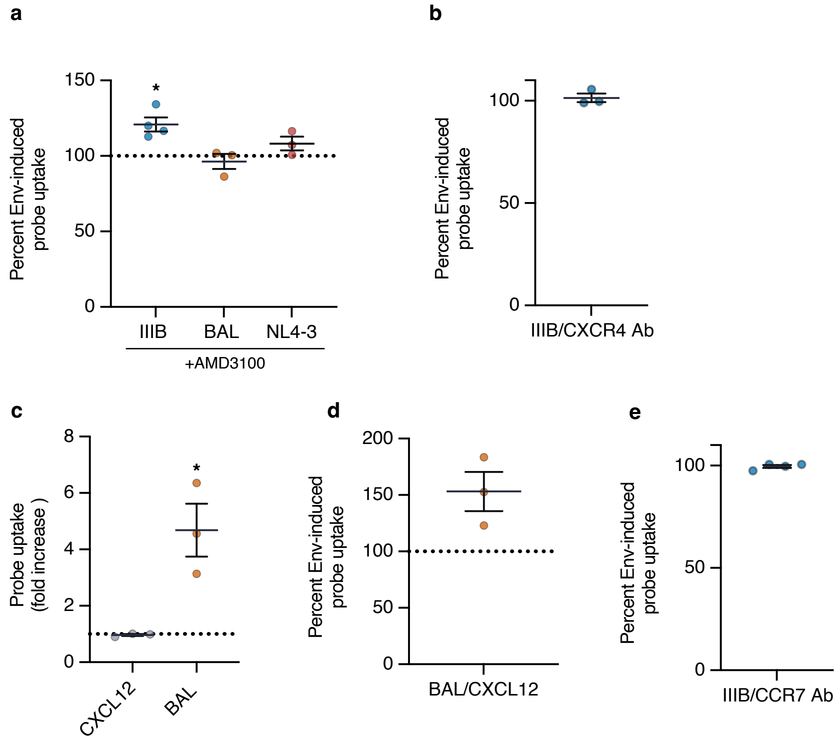

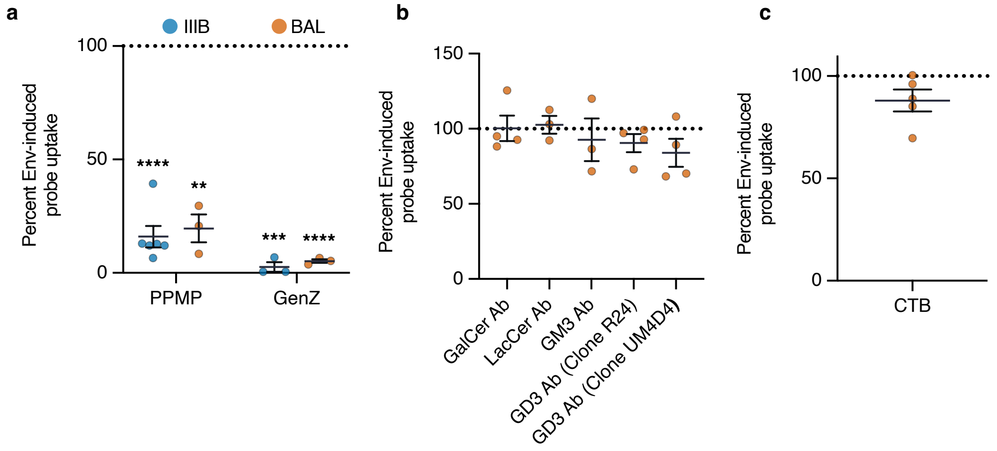
